# Modular, ultra-stable, and highly parallel protein force spectroscopy in magnetic tweezers using peptide linkers

**DOI:** 10.1101/491977

**Authors:** Achim Löf, Philipp U. Walker, Steffen M. Sedlak, Tobias Obser, Maria A. Brehm, Martin Benoit, Jan Lipfert

## Abstract

Single-molecule force spectroscopy has provided unprecedented insights into protein folding, force-regulation, and function. Here, we present a modular magnetic tweezers force spectroscopy approach that uses elastin-like polypeptide linkers to provide a high yield of protein tethers. Our approach extends protein force spectroscopy into the low force (<1 pN) regime and enables ultra-stable measurements on many molecules in parallel. We present (un-)folding data for the single protein domain ddFLN4 and for the large multi-domain dimeric protein von Willebrand factor (VWF) that is critically involved in primary hemostasis. The measurements reveal exponential force-dependencies of unfolding and refolding rates, directly resolve the stabilization of the VWF A2 domain by Ca^2+^, and discover transitions in the VWF C-domain stem at low forces that likely constitute the first steps of VWF activation *in vivo*. Our modular attachment approach will enable precise and multiplexed force spectroscopy measurements for a wide range of proteins in the physiologically relevant force regime.

## Introduction

Mechanical forces acting on proteins or ligand-receptor pairs are an integral part of many biological processes. Often the physiological function of proteins is critically regulated by force: examples include the mechano-activation of enzymes, force-regulated exposure of cryptic binding sites, and force-dependent unfolding and refolding of protein domains as “strain absorbers” to dissipate mechanical stress (*1*, *2*). A well-studied protein “strain absorber” is the fourth domain of the F-actin crosslinking filamin rod of *Dictyostelium discoideum* (ddFLN4), which exhibits extraordinarily fast refolding, facilitated by an intermediate state along the refolding pathway (*3*, *4*). Another remarkable example of physiological force regulation occurs in the large, multimeric glycoprotein von Willebrand factor (VWF) in the vasculature. VWF’s hemostatic function is regulated by increased hydrodynamic forces occurring upon blood vessel injury. Activation of VWF relies on a complex interplay of force-induced conformational changes both of single domains and of the large-scale protein conformation (*5*–*7*), while down-regulation of VWF is based on mechano-enzymatic cleavage at a cryptic binding site only accessible upon unfolding of VWF’s A2 domain (*8*). While many of the individual transitions in VWF have been probed in detail, the overall picture of how full-length VWF reacts to external forces in the blood stream remains incomplete. Since hydrodynamic peak forces grow as the square of the contour length (*5*, *8*), transitions that release contour length at low forces are expected to be particularly relevant for VWF’s physiological function as they will set of a cascade of increasing forces that trigger additional transitions with further contour length release. Recent work using AFM imaging has suggested transitions in the VWF C-domain stem that, however, could not be detected in AFM-based force spectroscopy, due to its limited force resolution (*9*, *10*).

Most insights into the mechanical properties and regulation of proteins and their complexes at the single-molecule level have been obtained from force spectroscopy experiments using atomic force microscopes (AFM) or optical tweezers (OT). While AFM and OT force spectroscopy measurements have provided unprecedented insights, they also have important shortcomings (*11*). AFM measurements cannot resolve forces below ~10 pN; OT provide excellent spatio-temporal resolution even for forces down to ~1 pN, but are not capable of measuring many molecules in parallel (*11*). In addition, both AFM and OT intrinsically control position and not force, such that constant-force measurements require active feedback. Magnetic tweezers (MT) are a single-molecule force spectroscopy technique that can overcome these shortcomings. In MT, molecules of interest are tethered between a surface and superparamagnetic beads (*11*–*13*) (**Supplementary Fig. S1**). External magnetic fields exert precisely controlled forces (*14*) in the range of ~0.01-100 pN and previous work has demonstrated camera-based tracking for ~10s-100s of nucleic acid-tethered beads simultaneously (*15*–*17*) in (*x*,*y*,*z*) with ~nm-spatial resolution, and, recently, also up to ≤ ms-temporal resolution (*18*–*20*). MT naturally operate in constant force mode, i.e. the applied force is constant during the measurement (to within 0.01%; **Supplementary Fig. S2**), as long as the external magnetic field is not actively changed, with excellent sensitivity in particular at low forces. In addition, MT enable long-term, stable, and robust measurements and do not suffer from heating or photo-damage (*11*).

Despite these advantages, MT so far have mainly been employed to investigate nucleic acid tethers. A key challenge in applying MT to protein force spectroscopy remains to tether ~nm-sized proteins between much larger, ~μm-sized beads and the surface, while avoiding unspecific surface interactions and ideally with a large number of usable tethers in each field of view. Previous MT studies on proteins, therefore, mostly employed large protein constructs, often polyproteins with repeats of e.g. titin Ig or protein L domains (*21*–*24*). Current strategies for attaching proteins to the surface in MT are either based on antibodies (*25*–*29*) or His-tag Cu^2+^-NTA chemistry (*30*, *31*), or on covalent linkage, either of Halo-tag fusion proteins to a surface coated with Halo-tag amine ligands (*21*–*23*, *32*) or using the Spy-tag-SpyCatcher system (*24*, *33*). Non-covalent attachment has the disadvantage of limited force stability compared to covalent attachment. Attachment via fusion proteins without a specific linker potentially complicates the analysis due to unfolding and refolding transitions of the proteins used for attachment (e.g. the Halo-tag (*21*)) or as spacers (e.g. flanking titin Ig-domains (*24*, *33*)). In addition, attachment via fusion proteins appears to suffer from a low number of usable tethers, as so far there are no reports of multiplexed protein unfolding and refolding measurements.

Here, we present a versatile, modular protein attachment strategy for single-molecule MT force spectroscopy. Our tethering protocol uses an elastin-like polypeptide (ELP) linker (*34*) that ensures efficient attachment to the surface while minimizing unspecific interactions, both critical prerequisites for high-throughput parallel measurements. In our approach, the protein of interest requires only short (1 and 11 amino acids [aa]) peptide tags for coupling to the linker and bead, respectively, avoiding the need for large fusion proteins and providing a general attachment strategy that is independent of protein size. We demonstrate the versatility of our attachment strategy by applying it to a small, single protein domain, ddFLN4 (100 aa), and a very large, multi-domain protein, dimeric full-length VWF (≈4000 aa). For both proteins, we achieve a high yield of specific tethers, i.e. a large number of single-molecule tethers that exhibit characteristic unfolding and refolding signatures and can be measured in parallel in a single field of view. Our highly-parallel ultra-stable measurements of repeated unfolding and refolding resolve outstanding questions about the respective folding pathways and stabilities. In addition, we leverage the ability of our assay to apply constant forces over extended periods of time to many molecules in parallel to probe the stability of the biotin-streptavidin receptor-ligand system. We anticipate our tethering strategy to be applicable to a wide range of proteins, and, furthermore, expect it to be of immediate use for other parallel force spectroscopy techniques, such as single-molecule centrifugation (*35*, *36*) or acoustic force spectroscopy (*37*), extending their capabilities towards multiplexed protein force spectroscopy.

## Results

### Site-specific and efficient tethering of proteins with elastin-like polypeptide linkers

Our attachment strategy uses an unstructured ELP linker (*34*, *38*) with a contour length of ≈120 nm and functional groups at its termini that we utilize as spacer for immobilizing the protein of interest to the bottom glass slide of the flow cell and to reduce unspecific protein-surface (*34*) and bead-surface interactions (**Fig. 1A**). The ELP linker is attached to a glass slide functionalized with thiol-reactive maleimide groups via an N-terminal cysteine (see **Methods** for details of the coupling protocol). The ELP linker carries a C-terminal *LPETGG* motif that allows for site-specific and covalent ligation to the protein of interest via an N-terminal glycine residue in a reaction catalyzed (*39*) by the enzyme sortase A. For coupling to the bead, the protein of interest is further engineered to carry an 11-aa ybbR-tag (*40*) at its C-terminus that is covalently attached to coenzyme A-biotin in the sfp phosphopantetheinyl transferase reaction. Finally, the biotin-label forms a high-affinity non-covalent bond to streptavidin-functionalized beads. Our approach requires only short peptide tags on the protein of interest that can readily be introduced by standard molecular cloning methods and have been shown to be compatible with expression and folding of a large range of proteins (*39*–*43*).

**Fig. 1.**
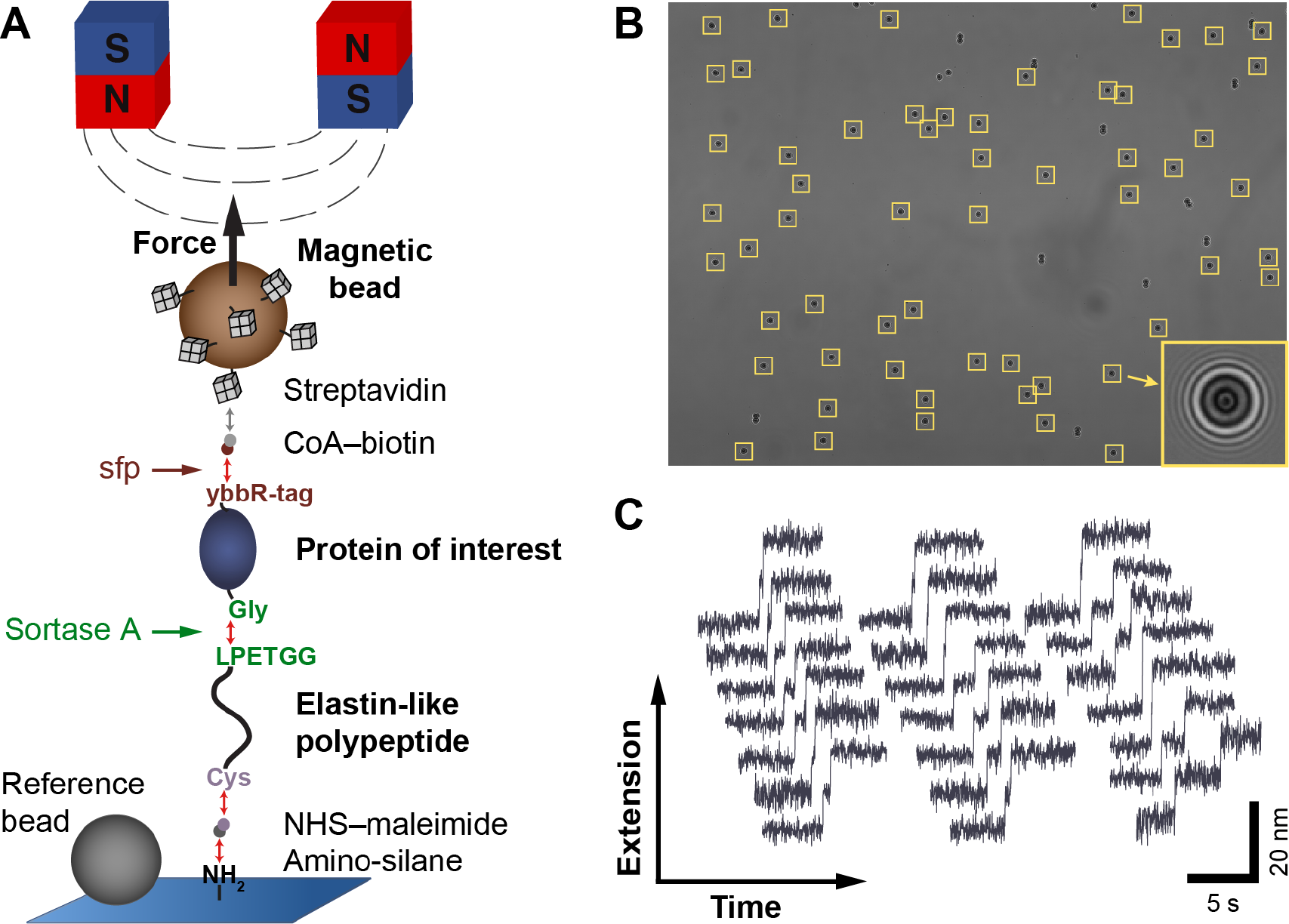
Attachment protocol for highly-parallel force spectroscopy on proteins in magnetic tweezers. (**A**) Schematic of the strategy for tethering a protein of interest between the bottom glass slide of the flow cell and a magnetic bead (not to scale). An ELP linker with a single cysteine at its N-terminus is coupled to the amino-silanized glass slide via a small-molecule NHS-maleimide cross-linker. After covalent coupling of coenzyme A (CoA)-biotin to the ybbr-tag at the C-terminus of the protein in a bulk reaction catalyzed by sfp phosphopantetheinyl transferase, the protein is covalently ligated to the ELP linker via one (or more) glycines at its N-terminus in a reaction mediated by sortase A, which selectively recognizes the C-terminal *LPETGG* motif of the ELP linker. Finally, a streptavidin-coated magnetic bead is bound to the biotinylated protein via the high-affinity biotin-streptavidin interaction. Red and grey double arrows indicate covalent and non-covalent bonds, respectively. Forces are exerted on the magnetic bead by permanent magnets positioned above the flow cell. Non-magnetic polystyrene beads baked onto the surface are used as reference beads for drift correction. (**B)** Representative field of view. Yellow boxes indicate approximately 60 beads marked for tracking. The enlarged image of one bead shows the diffraction ring pattern used for 3D bead tracking. (**C)** Example tether extension time traces showing the characteristic three-state unfolding pattern of ddFLN4. All traces shown were recorded in parallel from different beads within the same field of view at a constant force of 21 pN.

Here, we apply our tethering protocol to two very different protein systems: the small ddFLN4 domain and large full-length dimeric VWF. We obtained comparable and efficient tethering of beads with a large number of specific, single-molecule tethers in both cases. Typically, in a single field of view (≈440 × 330 μm^2^) of our MT setup (**Fig. 1B**; see **Methods** and **Supplementary Figs. S1** and **S2** for details on the setup) 50 to 100 tethered beads are tracked in parallel, of which 30 to 50 tethers exhibit characteristic unfolding and refolding signatures (**Fig. 1C**). The beads that do not show characteristic signatures are likely anchored to the surface by multiple protein tethers, since in control measurements without the protein of interest added, there is essentially no unspecific binding of beads to the surface (0-1 beads per field of view). The fraction of specific tethers attached via a single protein can be increased by decreasing the density of proteins immobilized on the surface. However, decreasing the protein concentration will also result in a decrease of the number of single-tethered beads. Optimizing our conditions, we achieved fractions of up to ≈60% specific, single-protein tethers, while still obtaining a large number of tethered beads at the same time. The most efficient flow cell exhibited 50 specific out of 85 beads within the single field of view measured.

### Three-state folding and unfolding of ddFLN4

We first applied our tethering protocol to the Ig-fold ddFLN4 domain (**Fig. 2A**), which exhibits a characteristic three-state unfolding pattern that has been extensively studied in AFM experiments (*3*, *4*) and is routinely employed as a molecular fingerprint in AFM force spectroscopy experiments (*44*–*46*). To characterize unfolding and (re-)folding in our MT assay, we recorded time traces of tether extension under different, constant forces. In a typical measurement (**Fig. 2B**), the force is increased from an initial low value (0.5 pN) that allows for (re-)folding, to a high value (25 pN in **Fig. 2B**) that promotes unfolding, and subsequently decreased to a moderate value (6.5 and 7.5 pN in **Fig. 2B**) to directly monitor refolding. Subsequently, this cycle is repeated multiple times with variable force levels to collect statistics. Unfolding and refolding of ddFLN4 were observed as clear double-steps in the traces, i.e., as an increase or decrease of the tether extension in two separate steps that we interpret as transition between the native (N) and intermediate (I) and between the intermediate and unfolded (U) states, respectively (**Fig. 2B**, insets). We analyzed the changes in extension for the transitions N↔I and I↔U as well as for the full transition N↔U for many different clamped forces (**Fig. 2C**). The resulting force-extension profiles are well described by fits of the worm-like chain (WLC) model with a fixed persistence length of 0.5 nm, in accordance with a previous AFM study (*4*), yielding contour length values (mean ± SD) of 15.0 ± 0.1 nm, 18.3 ± 0.1 nm, and 31.9 ± 0.2 nm, in excellent agreement with values reported from AFM (*3*, *4*).

**Fig. 2.**
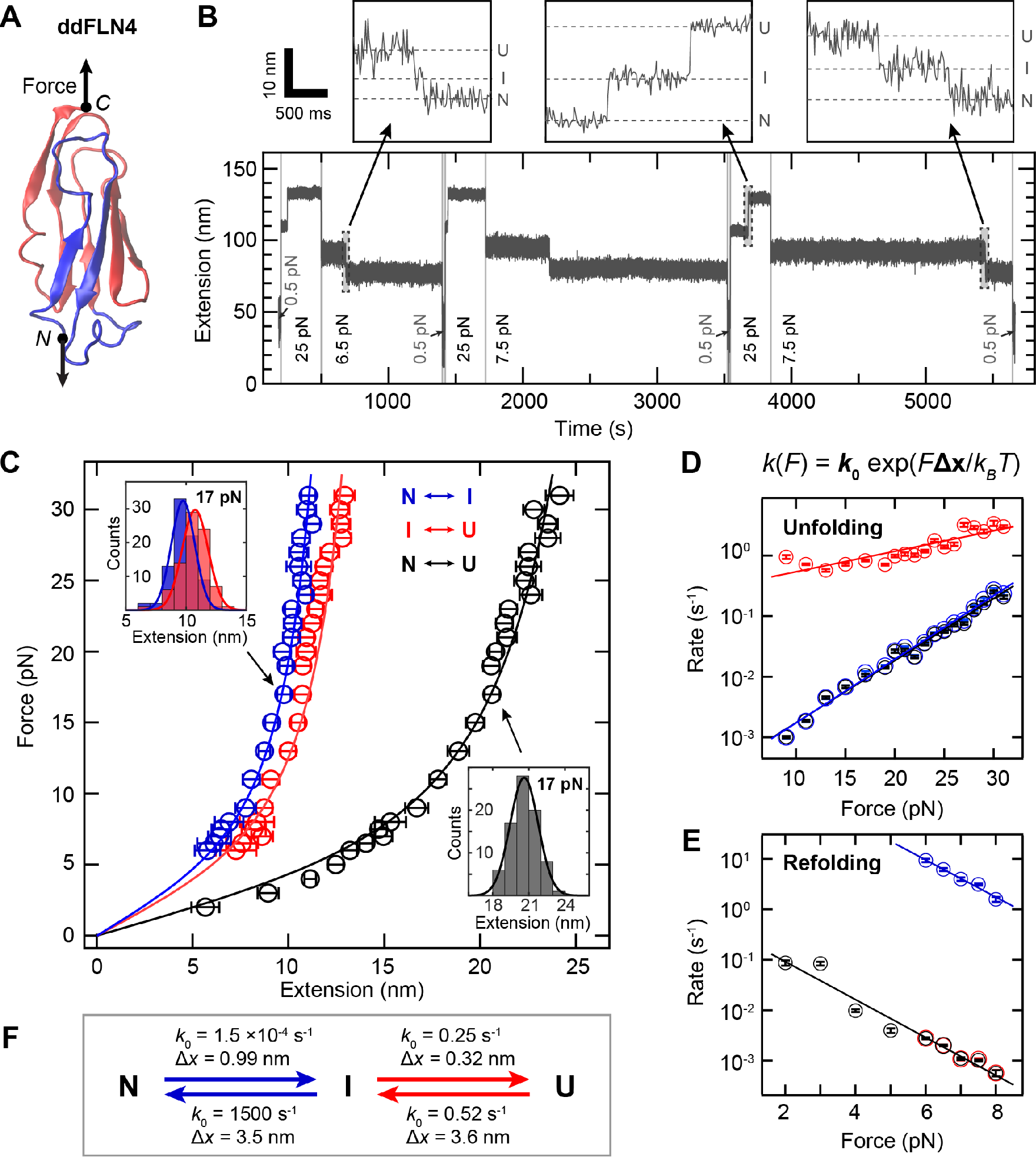
Three-state folding and unfolding of ddFLN4 at constant force. **(A)** Structure of ddFLN4 (PDB: 1KSR(*67*), rendered using VMD(*68*)), with strands A and B rendered in blue and strands C-G, forming the structured portion of the intermediate state, in red. Arrows indicate the direction of force acting on ddFLN4’s termini during MT experiments. **(B)** Extension vs. time trace of a ddFLN4 tether subjected to alternating intervals of high force (here 25 pN) allowing for ddFLN4 unfolding, intermediate force (here 6.5 or 7.5 pN) allowing for direct observation of refolding, and low force (0.5 pN) to ensure refolding before the next cycle. Unfolding and refolding of ddFLN4 via the mandatory intermediate state are observed as upward or downward double-steps in the traces, respectively. Dashed lines in insets indicate extension levels corresponding to the native (N), intermediate (I) and unfolded (U) states, respectively. **(C)** Force-extension profiles of ddFLN4 for the transitions N㆔I (blue) and I㆔U (red), and for full (un)folding N㆔U (black). Data points are obtained by Gaussian fits to step extension histograms (insets) at each constant force. Data points above 8 pN are from unfolding (based on 68-131 events obtained from 27-36 independent tethers), data points up to 8 pN from refolding (54-159 events from 26-39 independent tethers). Error bars correspond to the FWHM of Gaussian fits, divided by the square root of counts. Lines are fits of the WLC model. **(D)** Rates of unfolding at different constant forces for the three transitions. Color code as in panel C. Error bars correspond to 95% confidence intervals of exponential fits to the fraction of observed events as a function of time (**Methods**, **Supplementary Fig. S4**). Lines are fits of a single-barrier kinetic model. **(E)** Rates of refolding at different constant forces. Color code, error bars and fits analogous to panel D. **(F)** Fitted rates at zero force *k*_0_ and distances to the transition state Δ*x* for the unfolding and refolding transitions as determined from the fits of a single-barrier kinetic model shown in panels D and E.

Our data are fully consistent with previous work that found unfolding of the ddFLN4 domain to proceed via a mandatory, short-lived intermediate state: In a first unfolding step, strands A and B (42 aa; blue in **Fig. 2A**) detach and unfold, with strands C-G (58 aa; red in **Fig. 2A**) forming a less stable intermediate state(*3*), which quickly unfolds in the second unfolding step. Folding of ddFLN4 was also suggested to proceed via an intermediate state that is most likely structurally identical or very similar to the intermediate populated during unfolding (*4*). In our data set, data from unfolding (data points >8 pN) and refolding (data points ≤8 pN) are well described by a single WLC curve, confirming that the intermediate states populated during unfolding and folding are structurally very similar or identical. Importantly, no other features except the double-steps originating from ddFLN4 were observed in the force range probed (**Supplementary Fig. S3**), showing that the other components of our tethering strategy do not interfere with the measurements.

Our force clamp measurements allowed us to directly determine the rates of all transitions (**Methods**, **Supplementary Fig. S4**). For unfolding (**Fig. 2D**), we observed the rate for the first transition, NμI, to increase with increasing force from ≈ 0.001 s^−1^ at 9 pN to ≈ 0.2 s^−1^ at 31 pN. We fitted the rates to a single-barrier kinetic model, in which the rate is given by *k*(*F*) = *k*_0_ exp(*F*·Δ*x*/*k*_B_*T*), where *F* is the applied force, *k*_0_ the rate at zero force, Δ*x* the distance to the transition state, *k*_B_ the Boltzmann constant and *T* the temperature (*47*). We find 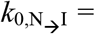 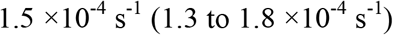 and 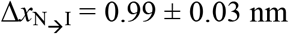 (mean ± SD for all rates and Δ*x* values). The measured rates for full unfolding N→U are essentially identical to those for the transition N→I (**Fig. 2D**), owing to the fact that the rates for the second transition, I→U 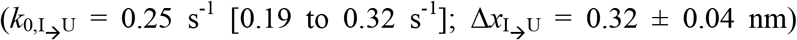, are at least one order of magnitude faster (**Fig. 2D**), implying that the transition N→I is the rate-limiting step for unfolding. The three-fold difference between 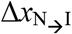 and 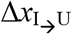 reflects that the difference between the rates N→I and I→U decreases with increasing force.

For refolding in the force range from 2 pN to 8 pN (**Fig. 2E**), the rates for the first substep U→I (*k*_0,U→I_ = 0.52 s^−1^ [0.34 to 0.79 s^−1^]; 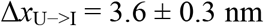) are again essentially identical to the rates for full refolding U→N (**Fig. 2E**) and rates for the second transition I→N (1500 s^−1^ [950 to 2500 s^−1^]; 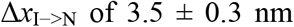) are at least three orders of magnitude higher compared to the first transition, implying that again the first transition, U→I, is rate-limiting (**Fig. 2F**). For forces below 6 pN, the intermediate state was too short-lived to be reliably detected, so that rates were determined separately only for forces ≥ 6 pN.

We compare our force clamp results to the rates at zero force reported previously from fits of a single barrier kinetic model to constant pulling speed AFM measurements (*4*). For unfolding, the rates for the second step *k*_0,I→U_ are in excellent agreement (0.25 and 0.33 s^−1^ in MT and AFM, respectively), yet the zero force rates for the first step *k*_0,N→I_ appear, in contrast, to differ significantly (1.5 ×10^−4^ *vs*. 0.28 s^−1^). However, in AFM measurements with extremely low pulling speeds of 1 nm/s a markedly higher mechanical strength of ddFLN4 has been observed and explained by switching to a second unfolding pathway (*48*). The reported zero-force rate (for full unfolding) from AFM of 1.4 ×10^−4^ s^−1^ is in excellent agreement with our results. Thus, our data support the existence of a second unfolding pathway and suggest that differences between the two pathways can be largely attributed to the first unfolding step N→I.

For refolding, a direct comparison is less straightforward, as refolding in AFM experiments has been measured at zero force and not under load (*4*). The rates obtained from MT and AFM differ significantly (*k*_0,U→I_: 0.52 *vs*. 55 s^−1^; *k*_0,I→N_: 1500 *vs*. 179 s^−1^), which might indicate different folding pathways in the presence and absence of force. Intriguingly, however, in both cases the same intermediate state appears to be populated during folding. Whereas the first step of folding –and thereby also full folding-is markedly slowed down by force, the second step of folding is almost 10-fold sped up, suggesting a pre-alignment of the structured portion of the intermediate state by force that allows for faster folding of strands A and B in the second folding step. Since ddFLN4 *in vivo* is positioned within actin-crosslinking filamin and under tensile load, it appears plausible that a force-induced pre-alignment of the intermediate state might play a physiological role.

### Ultra-stable equilibrium measurements of ddFLN4 unfolding and refolding

By determining the force at which the fitted rates for full unfolding and refolding (**Fig. 2D,E**; black lines) intersect, we predicted the equilibrium force at which the probabilities of ddFLN4 being in the unfolded and folded states are equal to be approximately 7.3 pN (**Fig. 3A**). We tested this prediction by measuring at a constant force of 7.5 pN close to the predicted equilibrium force. Since the predicted rates at equilibrium are only ~3 h^−1^ (**Fig. 3A**), we performed very long measurements (up to 55 h; **Fig. 3B**), harnessing the excellent force and drift stability of MT. We observed repeated transitions between the unfolded and folded states, with the system spending approximately half of the time in each of the two states, as expected for a measurement close to equilibrium. Examining the traces close to equilibrium in detail, we observe repeated transitions not only N↔U via the I state (**Fig. 3C**, left and middle trace), but also from the U and N states into the I state that return to the initial state (**Fig. 3C**, right trace), strongly suggesting that the same intermediate state is populated during unfolding and folding. Finally, we note that even for the very long measurements reported here, no significant change of ddFLN4’s force response over time was observed, indicating reliable, correct refolding of the domain without any hysteresis effects, as verified for a ≈10 h-long measurement comprising 35 unfolding and refolding cycles and for a long equilibrium measurement at constant force (**Supplementary Fig. S5**). The long-term stability combined with its very characteristic three-state unfolding signature make ddFLN4 an ideal fingerprint for the identification of single-molecule tethers.

**Fig. 3.**
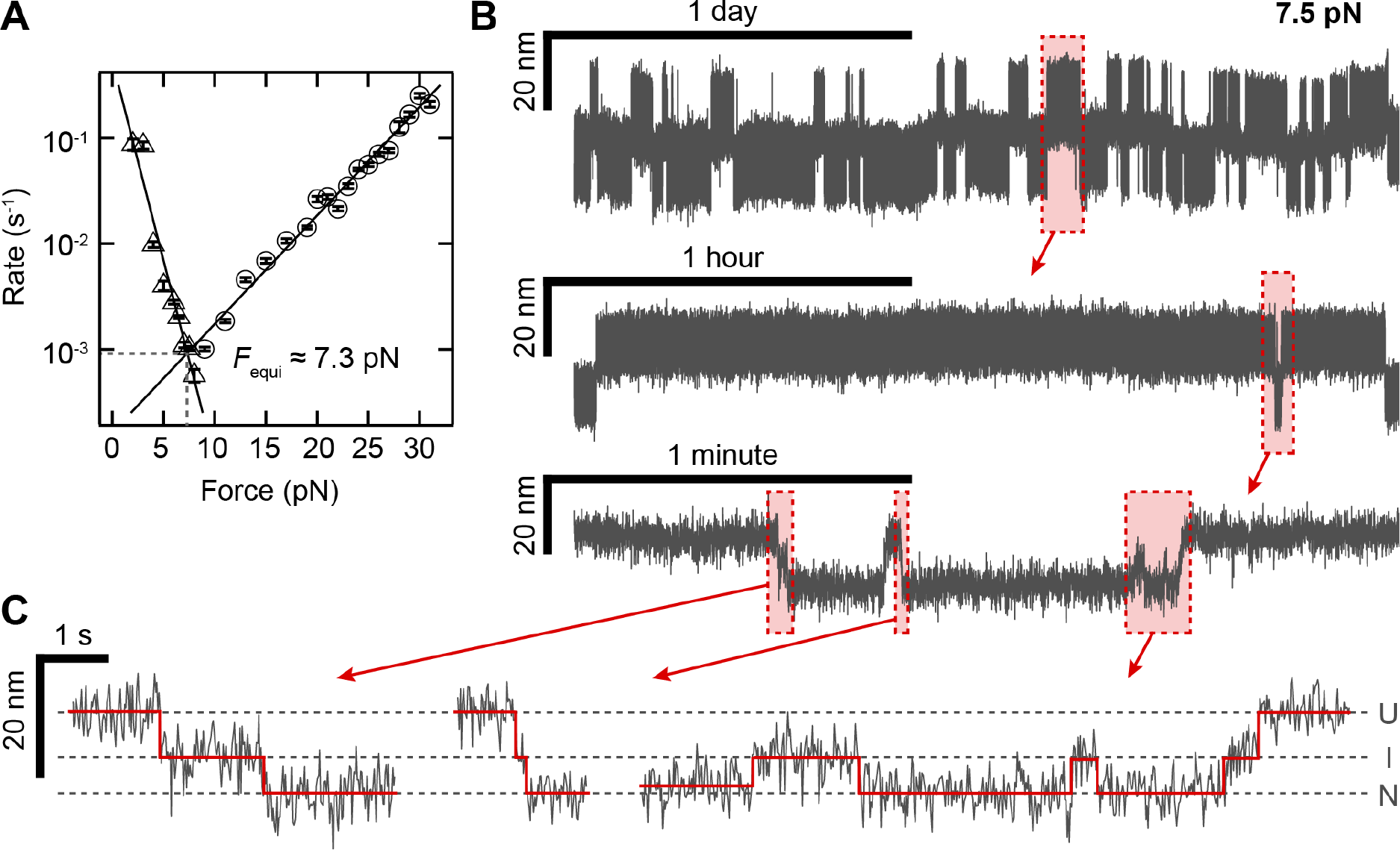
Long and stable ddFLN4 folding and unfolding traces close to equilibrium. **(A)** Force dependence of the rates for complete unfolding (circles) and refolding (triangles) as determined in Fig. 2d-e. The intersection of the linear fits predicts the equilibrium force *F*_equi_≈7.3 pN at which the probabilities of ddFLN4 being in the folded and unfolded state are equal. **(B)** 55 h-long extension *vs*. time trace of a ddFLN4 tether subjected to a constant force of 7.5 pN and zooms into indicated segments of the trace. **(C)** Zooms show not only full unfolding and folding transitions, but also transitions from the native to the intermediate state and back. Dashed lines indicate average extension levels corresponding to native (N), intermediate (I), and unfolded (U) state, respectively. Red lines indicate positions of transitions between states and extension levels in each state, as determined by the step-finding algorithm employed for analysis.

### Lifetime of biotin-streptavidin interactions for multi- and monovalent streptavidin

Having established ddFLN4 as an ideal fingerprint for the identification of specific singlemolecule tethers and having demonstrated the ability to apply constant forces over extended periods of time to multiple tethers in parallel, we utilized our assay to investigate not only protein folding and refolding, but also ligand-protein receptor interactions. As a proof-of-concept measurement, and to validate our tethering approach, we directly probed the stability of the high-affinity, non-covalent biotin-streptavidin interaction under constant force. Since all other linkages in our tethering protocol consist of mechanically stable covalent bonds, we used ddFLN4-tethered beads to apply different high forces (45-65 pN) to the biotin-streptavidin bond and monitored the time until bead rupture, only taking into account beads that showed the specific ddFLN4 unfolding signature in two short force plateaus of 25 pN at the beginning of the measurement. Importantly, the number of beads that ruptured already during these initial short plateaus was small (< 3.5%). For commercially available streptavidin-coated beads (Dynabeads M-270 Streptavidin, Invitrogen), we found the survival fraction to decay with time in a complex, multi-exponential fashion (**Fig. 4A**) for all forces probed, suggesting the existence of several populations of the biotin-streptavidin interaction with different lifetimes. To quantify the lifetimes involved, we fit the fastest and slowest decaying populations by linear regression to the logarithm of the first and last 20% of data points, respectively (lines in **Fig. 4A**). Over the studied force range, the lifetime of the fastest decaying population ranged from ≈100 s at 65 pN to ≈2100 s at 45 pN, whereas the lifetime of the slowest decaying population was ~50-fold higher, increasing from ≈5,000 s at 65 pN to ≈68,000 s at 45 pN (**Fig. 4C**). For both populations, the lifetime was found to increase exponentially with decreasing force (**Fig. 4C**). Already for a force of 20 pN, extrapolated lifetimes are well above a day and off-rates at zero force are in the range of 10^−7^ to 10^−8^ s^−1^, consistent with the fact that beads remained stably bound for hours or days in our force spectroscopy measurements at forces ≤ 20 pN (**Fig. 2,3, and 5**).

**Fig. 4.**
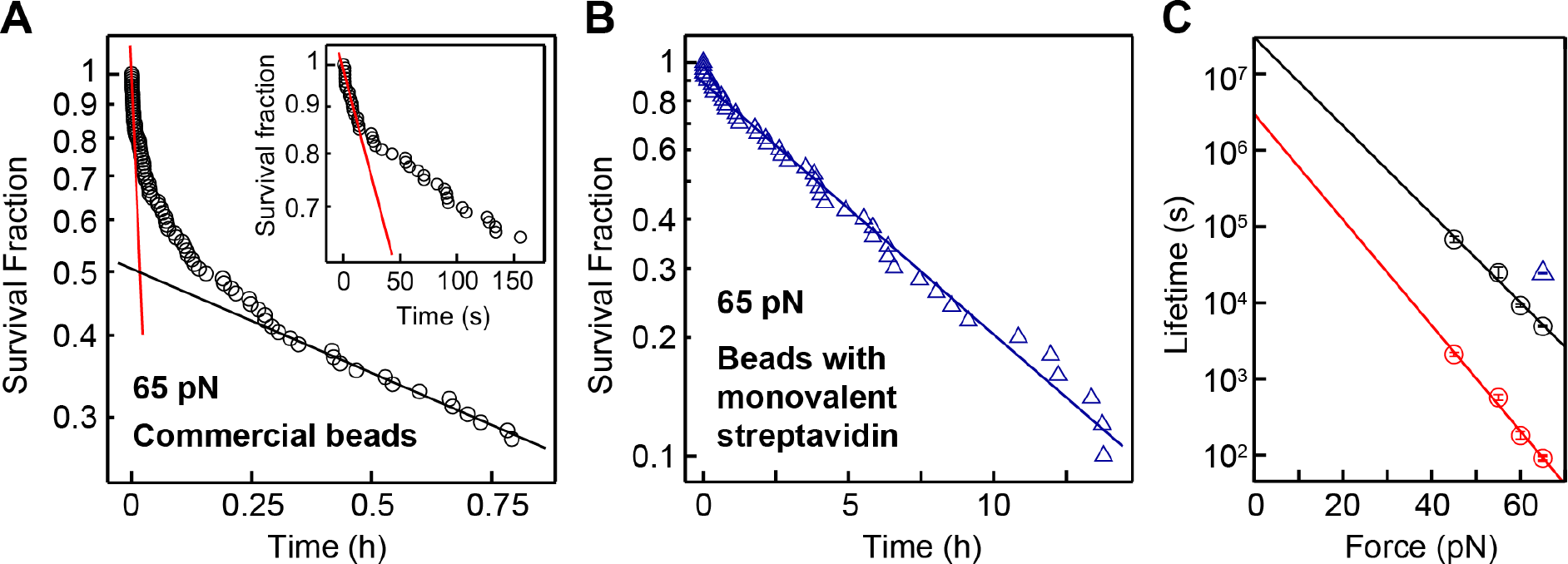
Bond lifetimes of beads functionalized with different streptavidin variants. **(A)** Survival fraction as a function of time for commercially available beads coated with tetravalent streptavidin (Dynabeads M-270 Streptavidin, Invitrogen), tethered by ddFLN4 and subjected to a constant force of 65 pN. The survival fraction decays in a complex, multiexponential fashion, suggesting the existence of several populations of biotin-streptavidin interactions with different lifetimes. Red and black lines are linear fits to the logarithm of the first and last 20% of data points, respectively, to estimate off-rates of the fastest-(inset) and slowest-decaying populations. **(B)** Survival fraction as a function of time for ddFLN4-tethered beads functionalized with a monovalent streptavidin variant, measured at 65 pN. The blue line is a linear fit to the natural logarithm of all data points. Note the markedly different scale of the time axis. **(C)** Estimated lifetime of the biotin-streptavidin interaction at different constant forces for the fastest-and slowest-decaying population of commercial beads with tetravalent streptavidin (red and black circles, respectively), and for beads with monovalent streptavidin (blue triangle). Error bars correspond to 1 SD; lines are fits of a single-barrier kinetic model. The number of measured, specifically tethered beads per condition was between 44 and 118, of which 35 to 86 ruptured during the duration of the measurement. The measurement duration was 15 h for beads with monovalent streptavidin and for the commercial beads 1 h at 65 pN, 3 h at 55 and 60 pN, and 15 h at 45 pN).

We hypothesized that the different populations and multi-exponential lifetimes for commercially available streptavidin-functionalized beads originate from the biotin-streptavidin complex being loaded with force in different geometries that result from the tetravalency of streptavidin (*46*). Indeed, for measurements with custom-made beads functionalized with a monovalent version of streptavidin (*46*) in a well-defined geometry using a C-terminal tag (*49*), the survival fraction was well described by a single-exponential decay (**Fig. 4B**). We chose immobilization of the monovalent streptavidin construct via the C-terminus of its functional subunit, as it has been recently demonstrated in AFM force spectroscopy measurements that the monovalent streptavidin-biotin complex can withstand markedly higher forces when loaded with force from the C-terminus as compared to pulling from the N-terminus (*49*). Indeed, we found the lifetime of the custom-made monovalent streptavidin beads to be 24,000 s (≈6.7 h) at 65 pN (**Fig. 4B**), similar to and even exceeding that of the 20% longest-lived commercially available beads (**Fig. 4C**).

### Force clamp measurements on full-length VWF dimers

Having demonstrated our attachment approach on a small well-characterized protein, we next applied it to large (≈500 kDa) dimeric constructs of full-length VWF. Dimers, the smallest repeating subunits of VWF multimers, consist of two multi-domain monomers that are C-terminally linked via disulfide bonds and have a contour length of ≈130 nm between their N-termini (*5*, *50*) (**Fig. 5A**). Since different peptide tags at the two N-termini are required for attaching dimers in the desired pulling geometry (**Fig. 5A**), we genetically engineered heterodimers consisting of two different monomers that are N-terminally modified with a ybbR-tag or a sortase motif GG, respectively (**Methods**). After tethering in the MT, we recorded time traces of VWF dimers with alternating plateaus of high force (**Fig. 5B**, 6-20 pN) and moderate force (**Fig. 5B**, 2-5 pN). In most cases, we observed two unfolding and two refolding steps in the recorded high and moderate force traces, respectively, with extension values matching the expected values for unfolding of the A2 domains (≈180 aa each) that were previously probed in isolation in OT (*8*, *51*). Observation of domain (un-)folding only for the two A2 domains is consistent with the prediction that all domains of VWF except A2 are protected against unfolding by long-range disulfide bonds (*50*, *52*) and with the results of recent AFM studies (*9*, *10*).

**Fig. 5.**
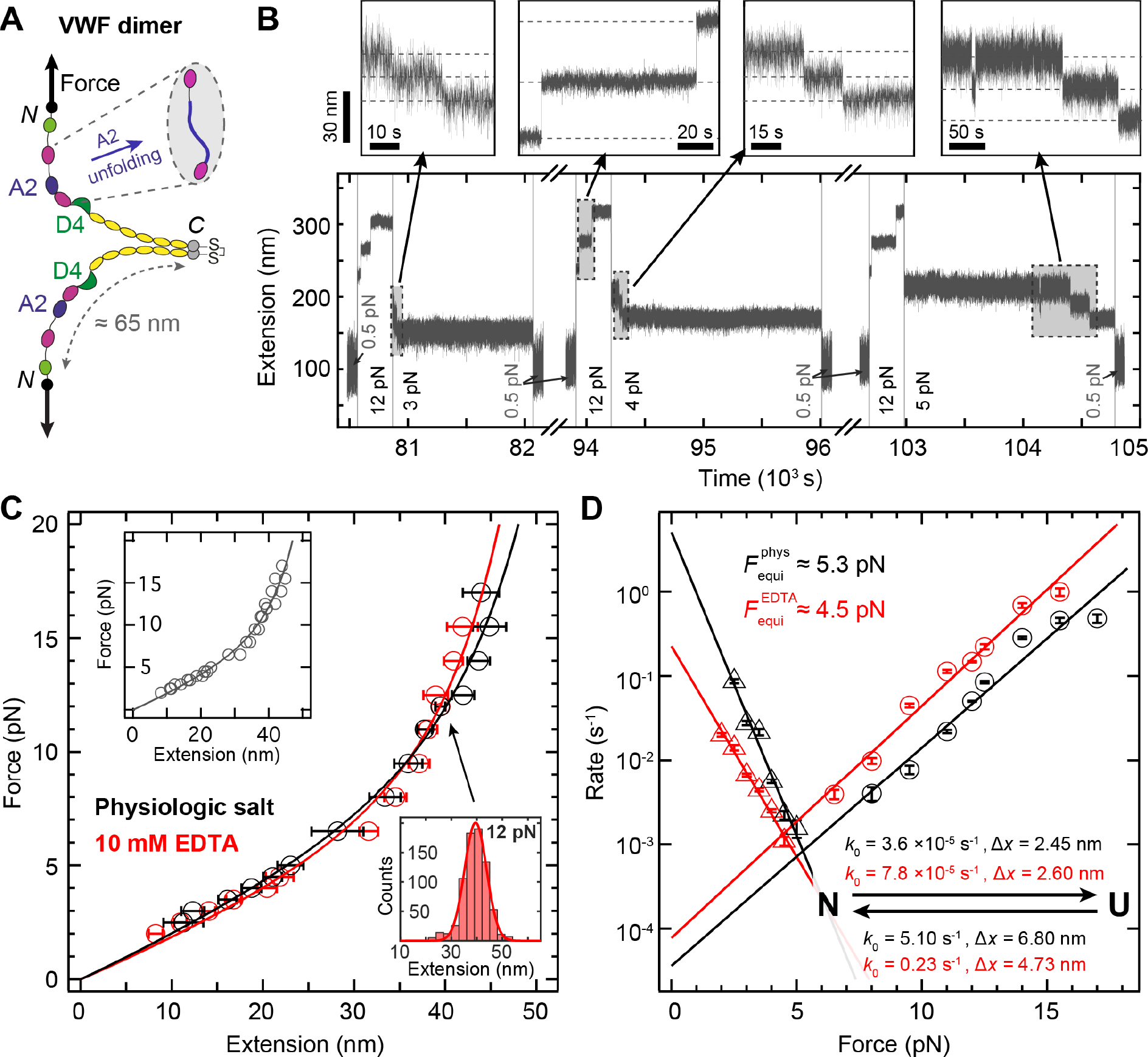
Folding and unfolding of A2 domains within VWF dimers. **(A)** Schematic structure of a VWF dimer, consisting of two ≈65 nm, multi-domain monomers C-terminally connected by disulfide bonds. The two A2 domains, which can unfold under force (inset), are shown in blue. Arrows indicate the direction of force acting on the two N-termini during MT experiments. **(B)** Segments from a ≈30-hour long extension *vs*. time trace of a VWF dimer tether subjected to alternating intervals of high force (here 12 pN), allowing for A2 unfolding, of intermediate force (here 3, 4 or 5 pN), allowing for direct observation of A2 refolding, and of low force (0.5 pN) to ensure refolding. Unfolding and refolding of the two A2 domains are observed as two independent positive or negative steps in the trace, respectively. Dashed lines in the insets indicate extension levels with none, one, or both of the A2 domains unfolded. **(C)** Force-extension curves of A2 (un)folding, in near-physiological buffer containing Ca^2+^ (black) and in buffer without Ca^2+^ and with 10 mM EDTA (red). Data points are obtained by Gaussian fits to step extension histograms (lower right inset) at each constant force. Data points above 5 pN are from unfolding (based on 62-632 and 40-747 events for the near-physiologic and EDTA case, respectively, obtained from 13-53 independent tethers), data points up to 5 pN are from refolding (41-120 and 49-158 events for the near-physiologic and EDTA case, respectively, obtained from 10-19 independent tethers). Error bars correspond to the FWHM of Gaussian fits, divided by the square root of counts. Lines are fits of the WLC model. Upper left inset shows a global WLC fit to all data points. **(D)** Rates of unfolding (circles) and refolding (triangles) at different constant forces for near-physiologic (black) and EDTA (red) buffer. Error bars correspond to 95% confidence intervals of exponential fits to the fraction of observed events as a function of time (**Methods**). Lines are fits of a single-barrier kinetic model, yielding rates at zero force *k*_0_ and distances to the transition state Δ*x* for unfolding and refolding as indicated.

In addition to the steps attributed to A2 unfolding and refolding, we less frequently also observed larger steps (**Supplementary Fig. S6**; 70-80 nm at ~11 pN), which we attribute to the dissociation of a strong intermonomer interaction mediated by the D4 domains that has recently been identified in AFM force measurements in approximately one-half of all VWF dimers under near-physiologic conditions (*9*, *10*). Consistent with their assignment to the D4-mediated intermonomer interaction, the large unfolding steps occur much less frequently in the absence of divalent ions, which have been shown to be critical for the intermonomer interaction (*9*, *10*), and are absent for mutant constructs lacking the D4 domain (delD4; **Supplementary Fig. S6,S7**). The dissociation of the intermonomer interaction was in some cases-after intermittent relaxation to a low force-observed repeatedly for the same molecule, implying reversibility of the interaction (**Supplementary Fig. S6**). Whereas in the constant pulling speed AFM measurements dissociation of this interaction had always occurred at much higher forces than –and therefore after– A2 unfolding, in our constant force measurements we observed dissociation of the D4-mediated intermonomer interaction in the same force range as A2 unfolding, suggesting a pronounced force-loading rate dependence for the intermonomer interaction. In fact, in the constant-force measurements we repeatedly observed dissociation of this interaction even before unfolding of one or both of the A2 domains (**Supplementary Fig. S6**). Importantly, this implicates a likely important role of the D4-mediated intermonomer interaction for regulation of VWF’s hemostatic activity at physiologically relevant forces in the bloodstream.

### Calcium binding stabilizes the VWF A2 domain

We next used our assay to elucidate the controversial impact of calcium on A2 stability (*51*, *53*, *54*). We performed measurements both in buffer mimicking the physiological pH and salt concentrations of the vasculature (‘near-physiologic’; pH 7.4, 150 mM NaCl, 1 mM MgCl_2_, and 1 mM CaCl_2_) and in buffer lacking divalent ions and supplemented with 10 mM EDTA. First, we analyzed the change in extension upon A2 unfolding and refolding for different constant forces. For both buffer conditions, the resulting force-extension profiles (**Fig. 5C**), combining data from unfolding (data points >6.5 pN) and from refolding (data points <5 pN), are well described by a single WLC curve. The WLC fits yielded values for contour length and persistence length of 75.0 nm and 0.42 nm (95% CI: 70.8–79.2 nm and 0.37–0.46 nm) for near physiological-buffer, and of 68.5 nm and 0.50 nm (62.7–74.3 nm and 0.41–0.58 nm) for the EDTA buffer, and thus show no significant difference, indicating that calcium has no effect on the extension of the unfolded state. A WLC fit to the combined data from both buffer conditions (inset in **Fig. 5C**) yielded contour and persistence length values of 71.9 nm and 0.45 nm (68.3–75.4 nm and 0.41–0.50 nm). The contour length increments determined from the MT measurements on full-length dimeric VWF are in excellent agreement with OT unfolding studies on isolated A2 domains (*8*, *51*, *53*), suggesting that complete A2 unfolding is not obstructed by the presence of other domains. Control measurements using the same attachment protocol and ddFLN4 under the same buffer conditions found no difference in the force-response for the different buffer conditions **Supplementary Fig. S8**.

Next, we studied the kinetics of A2 unfolding and refolding. In the case of unfolding, rates are approximately two-to fourfold higher for the EDTA buffer in the force range probed, 6.5-17 pN (**Fig. 5D**, circles). For both buffer conditions, rates increase exponentially with increasing force, with a slightly stronger dependence on force for the EDTA condition. Fitting a single-barrier kinetic model yielded values for the unfolding rate at zero force *k*_unf,0_ = 3.6 ×10^−5^ s^−1^ (1.8 to 7.1 ×10^−5^ s^−1^) and 7.8 ×10^−5^ s^−1^ (5.1 to 12 ×10^−5^ s^−1^) and distances to the transition state Δ*x*_unf_ = 2.45 ± 0.22 nm and 2.60 ± 0.15 nm in the presence and absence of Ca^2+^, respectively. The rates measured in our constant force assay are two orders of magnitude slower than the rates determined in near-physiological buffer in OT measurements on isolated A2 domains. While in principle this difference might indicate stabilization of A2 by neighboring domains, we deem it likely that it at least partially results from the transformation of rupture force distributions measured in the OT using the Dudko-Hummer-Szabo method (*55*), which is sensitive to the elastic response of employed flexible linkers. In our traces we did not observe any features that correspond to dissociation of potential interactions of A2 with neighboring domains. For refolding against external forces of 2-5 pN, rates are approximately two-to sixfold higher in the presence of Ca^2+^ (**Fig. 5D**, triangles) and decrease exponentially with force, with a more pronounced force dependence in the presence of Ca^2+^, which is reflected by the higher value of Δ*x* of 6.80 ± 0.56 nm compared to 4.73 ± 0.26 nm in the absence of Ca^2+^. The refolding rate at zero force in the presence of Ca^2+^ *k*_ref,0_ = 5.1 s^−1^ (2.9 to 8.7 s^−1^) is 20-fold higher than in the absence of Ca^2+^, *k*_ref,0_ = 0.23 s^−1^ (0.18 to 0.28 s^−1^), indicating that calcium substantially speeds up folding of A2.

Taken together, our results demonstrate that A2 is stabilized by the presence of Ca^2+^ by increasing the refolding rate and stabilizing against unfolding compared to the conditions without Ca^2+^. The observed increases in the refolding rates in our experiments are in quantitative agreement with a previous report using OT on isolated A2 domains (*51*). Importantly, we directly observe refolding under mechanical load even in the absence of Ca^2+^ (**Supplementary Fig. S9**), in contrast to a previous study (*53*). The role of Ca^2+^ in the stabilization against unfolding is controversial: We observe a modest reduction in the unfolding rate by Ca^2+^, which is consistent with the low-force data found in one OT study (*51*), which, however, reported no statistically significant change in the unfolding rate with and without Ca^2+^ overall, possibly as their assay might have lacked the sensitivity to resolve small differences. In contrast, we find no evidence for a long-lived intermediate in the unfolding pathway in the presence of Ca^2+^ that was reported by another study using OT (*53*). Finally, we occasionally observed tethers that only showed the unfolding and refolding signal of one A2 domain (**Supplementary Fig. S10**). In such tethers, refolding of one A2 domain may be inhibited due to cis-trans isomerization of a cis-proline, as reported in a previous OT study (*8*).

### Transitions in the VWF stem at low forces

AFM imaging (*9*, *10*) and electron microscopy (*56*) suggest that the VWF stem consisting of six C-domains can open and close in a zipper-like fashion (**Fig. 6A,B**). However, transitions of the VWF stem have not been observed directly. To probe for interactions in the VWF stem, we subjected dimeric VWF tethers that had shown the characteristic A2 unfolding pattern to low constant forces. At forces of ~1 pN, we observed repeated, reversible transitions with a maximum contour length increase of ~50-60 nm that is consistent with fully unzipping and rezipping the VWF stem (**Fig. 6C**). Increasing force in the range 0.6-1.4 pN systematically shifted the population towards higher tether extensions, which we interpret as less compact “unzipped” conformations of the VWF stem. Importantly, such transitions were never observed for ddFLN4-tethered beads. In addition, the large change in extension at very low forces makes it appear highly unlikely that the observed transitions originate from domain unfolding events (for comparison, a tether extension of < 5 nm would be expected for unfolding of an A2 domain at a force of ~1 pN, due to the WLC stretching behavior of the unfolded protein chain).

**Fig. 6.**
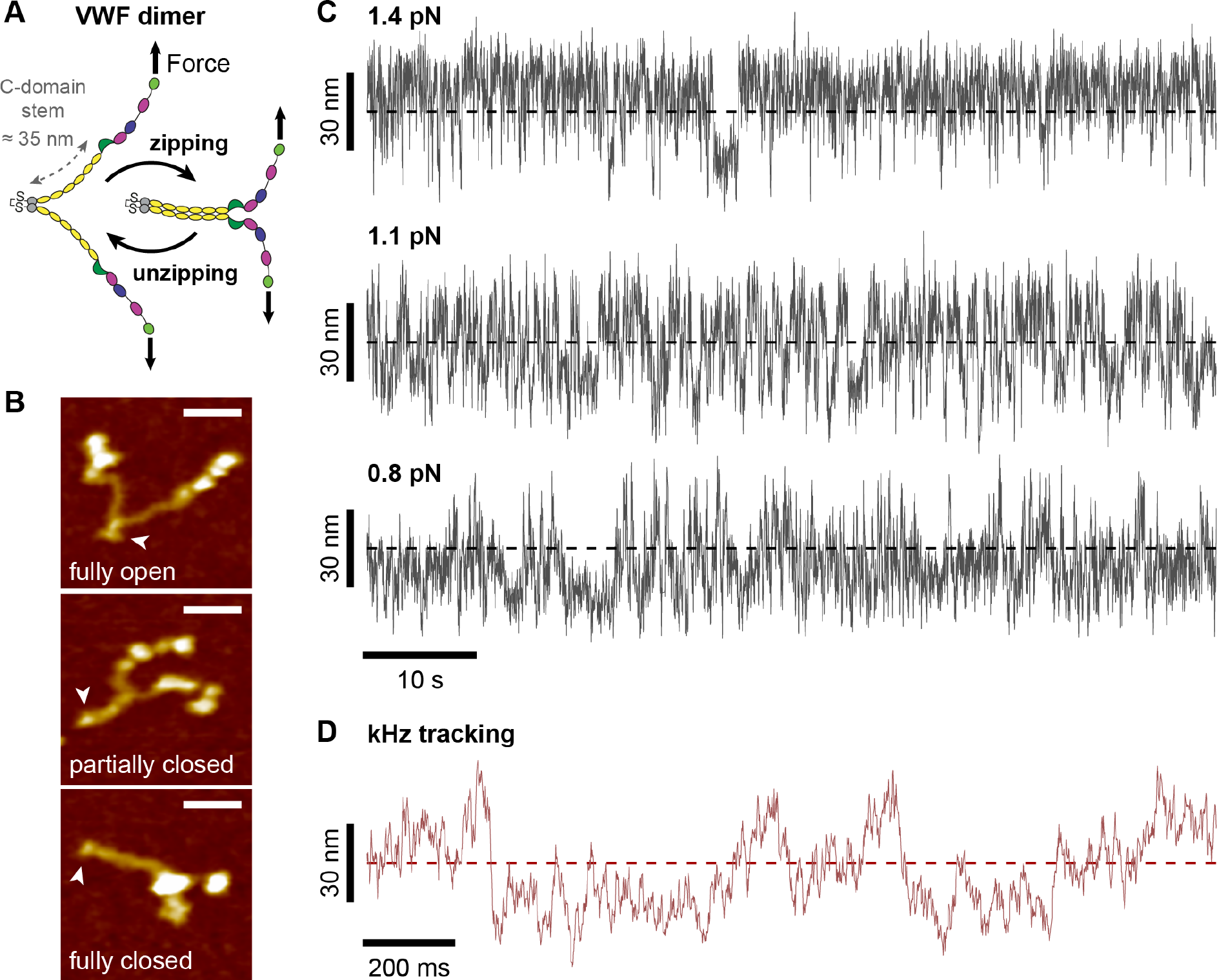
Low-force unzipping and zipping of the C-domain stem in VWF dimers. **(A)** Schematic of closing and opening of the stem (domains C1-C6, yellow) of VWF dimers in a zipper-like fashion due to interactions between the C-domains. **(B)** AFM images of single VWF dimers adsorbed onto a surface under near-physiologic buffer conditions. Arrowheads indicate C-terminal ends of dimers, where the two constituting monomers are linked. In thermal equilibrium and in the absence of force, dimer stems exhibit conformations ranging from fully open to fully closed. It should be noted that in approximately one-half of all dimers, the stem region is firmly closed by the D4-mediated intermonomer interaction (*9*, *10*). Scale bars are 20 nm; height range of color scale is 2.4 nm. **(C)** Extension *vs*. time traces of the same VWF dimer tether subjected to low forces. Fast reversible transitions between a maximum and minimum value of extension, approximately 60 nm apart, are observed that we attribute to closing and opening of the C-domain stem. Dashed lines indicate the midpoint between the two extreme values of extension. At a force of ≈1.1 pN, the system spends approximately half of the time above and below the midpoint. Traces are raw data recorded at 58 Hz. **(D)** Segment of an extension vs. time trace of the same tether shown in panel c, at 1.3 pN, recorded at an acquisition rate of 1 kHz. The measurements with high temporal resolution confirm that the observed transitions are not jumps between two discrete extension levels, but rather gradual transitions with several intermediate extension levels, in line with a zipper-like closing and opening of the stem with several pairs of interacting C-domains.

The observed transitions occurred between multiple levels and featured more than two distinct states, as expected from the observation by AFM imaging that multiple C-domain interactions can contribute to the opening and closing of the stem. However, a reliable assignment of the transitions will require additional analysis and measurements under a range of solution conditions and using VWF mutants. Using high-speed bead tracking (1 kHz; **Fig. 6D**), we observed transitions to occur on time scales ≤ 1 s. Our observations are consistent with the predicted (*9*) occurrence of unzipping transition in the VWF stem at forces of ~1 pN, close to the value for the onset of VWF multimer elongation predicted by Brownian hydrodynamics simulations (*57*). Molecular interactions that break and release contour length at such low forces are expected to be particularly relevant for VWF’s physiological function as these are likely the first interactions to open under shear flow and to set off a cascade of increased contour length and increased force, since hydrodynamic peak forces grow as the square of the contour length (*5*, *8*).

## Discussion

We have introduced a novel approach for single-molecule force spectroscopy measurements on proteins using MT. Our protocol enables multiplexed measurements through high-yield, ultra-stable tethering of proteins requiring only short peptide tags. As a proof-of-concept measurement, we probed the three-state unfolding and refolding of ddFLN4. Our measurements at constant force are overall in excellent agreement with constant pulling speed AFM experiments and confirm the existence of a low-loading rate pathway for unfolding (*4*, *48*). Applying our method to the large, force-regulated protein VWF confirms several structural transitions previously observed in AFM and OT force-spectroscopy measurements and reveals how Ca^2+^ stabilizes the A2 domain. In addition, our measurements reveal transitions in the VWF stem at low forces of ≈1 pN, which likely constitute critical first steps in the stretch response of VWF under physiological shear flow.

Using the ability of our assay to apply constant forces over long periods of time to multiple tethers in parallel, we probed the stability of streptavidin-biotin bonds, a widely used ligand-receptor system. We found that commercially available, multivalent streptavidin (far) exceeds the requirements of typical constant force force-spectroscopy measurements (with lifetimes ≥ 1 day at forces ≤ 20 pN), but has a multi-exponential lifetime distribution. Monovalent, site-specifically attached streptavidin, in contrast, exhibits a single-exponential lifetime distribution with extremely high force stability, making it an attractive approach for force spectroscopy on systems that require high forces over extended periods of time. Ultimately, one could also replace the biotin-streptavidin bond with a covalent linkage to even further enhance the force and also chemical stability of the attachment protocol.

In conclusion, we have demonstrated the versatility and power of a new approach for singlemolecule protein force spectroscopy measurements using MT. Our method provides a high yield of ultra-stable specific single-molecule tethers that can be probed in parallel at constant forces over extended periods of time. Given ongoing improvements in camera technology, we expect that the number of protein tethers that can be measured in parallel will further increase by at least an order of magnitude in the near future. In addition, we anticipate that our tethering strategy will enable multiplexed protein force spectroscopy using other singlemolecule methods such as acoustic and centrifugal force spectroscopy as well. Since the approach is modular and only requires minimal modifications to the protein of interest, we anticipate it to be applicable to a wide range of proteins. We expect MT force spectroscopy to in particular give access to the physiologically relevant low force (< 1 pN) regime and to provide a wealth of novel insights into the mechanics and force-regulation of proteins.

## Materials and Methods

### Preparation of ddFLN4 constructs

Recombinant ddFLN4 expressed in *E.coli* (with the internal cysteine at position 18 mutated to serine) was a kind gift from Lukas Milles (LMU Munich). At its C-terminus, the ddFLN4 construct possesses a polyhistidine-tag for purification and a ybbR-tag (*40*). At its N-terminus, the construct has a short linker sequence (*MGTGSGSGSGSAGTGSG*) with the terminal methionine followed by a single glycine. Due to efficient cleavage of the methionine by *E.coli* methionine aminopeptidases, the glycine is expected to be available for sortase-catalyzed ligation.

The ddFLN4 gene was synthesized codon-optimized for expression in *E.coli* as a linear DNA fragment (GeneArt – ThermoFisher Scientific, Regensburg, Germany), and inserted into pET28a Vectors via Gibson assembly (*58*) (New England Biolabs, Frankfurt, Germany). Protein expression in *E.coli NiCo21 (DE3)* (New England Biolabs) and purification via the polyhistidine-tag were carried out as previously described in detail(*45*).

### Preparation of hetero-bifunctional VWF dimer constructs

For preparation of hetero-bifunctional VWF dimers two different types of monomers were coexpressed, which at their N –termini-subsequent to a required signal peptide– possess either a ybbR-tag (*40*) or an N-terminal strep-tag II for purification (*59*), followed by a tobacco etch virus (TEV) protease cleavage site (*60*) and the sortase motif *GG* (*39*). The TEV site serves two purposes: first, to remove the strep-tag after purification, as it might otherwise interact with Streptavidin on the magnetic beads during measurements, and second, to free the sortase motif *GG*, which must be located terminally for the sortase reaction. Both monomer constructs lack the VWF pro-peptide (domains D1 and D2) in order to abolish linkage of dimers into larger multimers. For delD4 dimers, additionally the D4 domain is deleted in both monomers. For AFM images shown in Fig. 5, dimeric VWF constructs consisting of two identical monomers, possessing a Strep-tag at their N-termini, were used.

Plasmid construction was carried out analogously to a procedure previously described(*9*). For expression, 2·10^6^ HEK 293 cells in a 75 cm^2^ flask (DSMZ, Braunschweig, Germany) were transfected in Dulbecco’s modified Eagle’s medium (Life Technologies, Darmstadt, Germany) containing 10% fetal bovine serum (Life Technologies), 2 μg of each of the two plasmids, and 15 μl Lipofectamine 2000 (Life Technologies). 24 h after transfection, cells were transferred into selection medium containing 500 μg/ml G418 (Invivogen, Toulouse, France) and 250 μg/ml Hygromycin B (Invivogen). After 2–3 weeks, the polyclonal cell culture was seeded for expression. After 72 h of cell growth, the medium was exchanged against OPTIPRO-SFM (Life Technologies) for serum-free collection of secreted recombinant VWF. The culture supernatant was collected after 72 h and concentrated using Amicon Ultra-15 MWCO 100 kDa (Merck, Schwalbach, Germany).

Dimeric constructs were purified via a HiTrap StrepTrap affinity chromatography column (GE Healthcare) using the AEKTA Explorer system (GE Healthcare). As running buffer, 20 mM 4-(2-hydroxyethyl)-1-piperazineethanesulfonic acid (HEPES), 150 mM NaCl, 1 mM MgCl_2_, 1 mM CaCl_2_, pH 7.4, was used. Elution buffer additionally contained 2.5 mM d-desthiobiotin. Eluates were buffer exchanged (to the running buffer) and concentrated by centrifuge filtration using Amicon Ultra MWCO 100 kDa (Merck Millipore). All purified VWF dimers were further inspected by AFM imaging and showed no structural differences as compared to dimeric VWF constructs with different peptide tags or without tags used in previous studies (*9*, *10*).

### Preparation of ELP linkers

Recombinant ELP linkers expressed in *E.coli NiCo21 (DE3)* were a kind gift from Wolfgang Ott (LMU Munich). The ≈300 aa ELP linker with a contour length of ≈120 nm used in this study has the sequence [(*VPGEG*)-(*VPGVG*)_4_-(*VPGAG*)_2_-(*VPGGG*)_2_-(*VPGEG*)]_6_ and possesses a single N-terminal cysteine and the C-terminal sortase recognition motif *LPETGG*. Cloning, expression and purification have been described (*34*, *38*), and can be performed using standard procedures for the production of recombinant proteins. Plasmids are provided at Addgene by Ott et al. (Addgene accession number 90472 for the ELP linker used in this study).

### Attachment chemistry and flow cell preparation

Functionalization of glass slides with the ELP linkers described above followed the protocol by Ott et al. (*34*). Glass slides were first silanized with 3-(aminopropyl)dimethylethoxysilane (APDMES, ABCR GmbH, Karlsruhe, Germany), and then coated with 10 mM of a sulfosuccinimidyl 4-(N-maleimidomethyl)cyclohexane-1-carboxylate cross-linker with a negligible contour length of 0.83 nm (Sulfo-SMCC, Thermo Fisher Scientific Inc.), dissolved in 50 mM HEPES, pH 7.5. Subsequently, ELP linkers were linked to the thiol-reactive maleimide groups via the single cysteine at their N-terminus in coupling buffer consisting of 50 mM sodium phosphate, 50 mM NaCl, 10 mM EDTA, pH 7.2. Afterwards, 10 mM L-cysteine dissolved in coupling buffer were added to saturate potentially remaining unreacted maleimide groups. Finally, non-magnetic polystyrene beads (Polybead Microspheres 3 μm; Polysciences GmbH, Hirschberg, Germany) dissolved in ethanol were baked onto the slides at ≈70 °C for ≈5 min for use as reference beads. After each step, slides were extensively rinsed with ultrapure water. Flow cells were assembled from an ELP-functionalized cover slip as the bottom surface and a non-functionalized cover slip with two small holes for inlet and outlet as the top, with a layer of cut-out parafilm (Pechiney Plastic Packaging Inc., Chicago, IL) as a spacer to form a (~4 mm wide and 50 mm long) flow channel. Flow cells were assembled by heating on a hot plate to ≈70 °C for ≈2 min. Assembled flow cells can be stored under ambient conditions for weeks.

Prior to experiments, the flow cells were incubated with 1% casein solution (Sigma-Aldrich) for 1 h and afterwards flushed with 1 ml (approximately 20 flow cell volumes) of buffer (20 mM HEPES, 150 mM NaCl, 1 mM MgCl_2_, 1 mM CaCl_2_, pH 7.4). CoA-biotin (New England Biolabs) was coupled to the ybbR-tag on the protein of interest in a bulk reaction in the presence of 5 μM sfp phosphopantetheinyl transferase and 10 mM MgCl_2_ at 37 °C for 60 min. In the case of VWF, subsequently TEV protease was added to a final concentration of approximately 25 μM and incubated for 30-60 min. Dithiothreitol (DTT) present in the storage buffer of TEV protease was removed beforehand using desalting columns (Zeba Spin 40 K MWCO, Thermo Scientific Inc.). Afterwards, protein was diluted to a final concentration of approximately 10 nM (VWF dimers) or 25 nM (ddFLN4) in 20 mM HEPES, 150 mM NaCl, 1 mM MgCl_2_, 1 mM CaCl_2_, pH 7.4, and incubated in the flow cell in the presence of 1-2 μM sortase A for 30 min. Subsequently, the flow cell was flushed with 1 ml of buffer.

Magnetic beads –either Dynabeads M-270 streptavidin (Invitrogen) or beads functionalized with monovalent streptavidin (see below)– in measurement buffer containing 0. 1% (v/v) Tween-20 (Sigma-Aldrich) were incubated in the flow cell for 60 s, and unbound beads were flushed out with 2 ml of measurement buffer. All measurements were performed at room temperature (≈22 °C).

Starting with silanized glass slides, complete flow cell preparation takes less than 7 h. In addition, flow cells functionalized with ELP linkers, but not yet incubated with casein and protein, can be prepared in advance and stored at room temperature for weeks without loss of functionality. Starting with ELP-functionalized flow cells, measurements can be started within 120 min.

### Preparation of monovalent streptavidin

Tetrameric, but monovalent streptavidin (mSA) consisting of three mutant subunits deficient in biotin binding and one functional subunit, possessing at its C-terminus a polyhistidine-tag for purification and a single cysteine for site-specific immobilization, was prepared as described in detail by Sedlak et al. (*46*, *49*). In brief, functional and mutant subunits were cloned into pET vectors (Novagen, EMD Millipore, Billerica, USA) and separately expressed in *E.coli BL21(DE3)-CodonPlus* (Agilent Technologies, Santa Clara, USA). Resulting inclusion bodies were solubilized in 6 M guanidine hydrochloride. Functional and mutant subunits were then mixed at a 1:10 ratio prior to refolding and purification via the polyhistidine-tag, in order to ensure a 1:3 ratio of functional to non-functional subunits in the final tetrameric streptavidin construct.

### Site-specific, covalent immobilization of monovalent streptavidin on magnetic beads

Magnetic beads with surface amine groups (Dynabeads M-270 Amine, Invitrogen; these beads are otherwise identical to Dynabeads M-270 Streptavidin) were functionalized with 25 mM of 5-kDa NHS–polyethylene glycol (PEG)–maleimide linkers with reactive NHS and maleimide end groups (Rapp Polymere, Tübingen, Germany) in 50 mM HEPES, pH 7.5, and afterwards extensively washed first with DMSO and then with water. The mSA constructs possessing a single cysteine as described above were reduced with 5 mM TCEP bond breaker solution (Thermo Fisher) and afterwards buffer exchanged to coupling buffer using desalting columns (Zeba Spin 40 K MWCO, Thermo Scientific Inc.). Beads were then incubated with mSA in coupling buffer for 90 min and extensively washed with measurement buffer.

### Buffers

All measurements on ddFLN4 and measurements on VWF dimers under ‘near-physiologic’ conditions were performed in buffer containing 20 mM HEPES, 150 mM NaCl, 1 mM MgCl_2_, 1 mM CaCl_2_, 0.1% Tween-20, at pH 7.4. Measurements without calcium were performed in EDTA buffer containing 20 mM HEPES, 150 mM NaCl, 10 mM EDTA, 0.1% Tween-20, at pH 7.4. Before measurements in the absence of calcium, the flow cell was incubated with EDTA buffer for 2 h. Control measurements at acidic pH were performed in 20 mM sodium-acetate, 150 mM NaCl, 1 mM MgCl_2_, 1 mM CaCl_2_, 0.1% Tween-20, at pH 5.5.

#### Magnetic tweezers setup

Measurements were performed on a custom MT setup described by Walker et al. (*61*). A schematic and an image of the setup are given in **Supplementary Fig. S1**. The setup uses a pair of permanent magnets (5×5×5 mm^3^ each; W-05-N50-G, Supermagnete, Switzerland) in vertical configuration (*14*). The distance between magnets and flow cell (and, therefore, the force; **Supplementary Fig. S2**) is controlled by a DC-motor (M-126.PD2; PI Physikinstrumente, Germany). For illumination, an LED (69647, Lumitronix LED Technik GmbH, Germany) is used. Using a 40x oil immersion objective (UPLFLN 40x, Olympus, Japan) and a CMOS sensor camera with 4096×3072 pixels (12M Falcon2, Teledyne Dalsa, Canada), a large field of view of approximately 440 × 330 μm^2^ can be imaged at a frame rate of 58 Hz. For measurements with an acquisition rate of 1 kHz, a reduced field of view of 1792 × 280 pixels was used. Images are transferred to a frame grabber (PCIe 1433; National Instruments, Austin, TX) and analyzed with an open-source tracking software (*15*). The bead tracking accuracy of our setup was determined to be ≈0.6 nm in (*x*, *y*) and ≈1.5 nm in *z* direction, as determined by tracking non-magnetic polystyrene beads, with a diameter comparable to M270 beads (3 μm), after baking them onto the flow cell surface. For creating the look-up table required for tracking the bead positions in *z*, the objective is mounted on a piezo stage (Pifoc P-726.1CD, PI Physikinstrumente). Force calibration was performed as described by te Velthuis et al. (*62*) based on the fluctuations of long DNA tethers. The final force calibration, i.e. the dependence of the force applied to a bead on the distance between magnets and flow cell, is shown in **Supplementary Fig. S2**, together with an example trace showing the DNA B-S overstretching transition at the expected force of ≈65 pN. Importantly, for the small extension changes on the length scales of our protein tethers, the force stays constant to very good approximation, with the relative change in force due to tether stretching or protein (un-)folding being < 10^−4^ (**Supplementary Fig. S2**). We verified the uniformity of the magnetic field across the field of view and found the change in force across the full range of the field of view to be < 3% (**Supplementary Fig. S2**). The largest source of force uncertainty is the bead-to-bead variation, which we found to be on the order of ≤ 10% for the beads used in this study (**Supplementary Fig. S2**), in line with several previous reports (*14*, *63*, *64*).

#### AFM imaging

For AFM imaging, a dimeric VWF construct possessing a strep-tag at both N-termini was used. Preparation of substrates for AFM imaging was performed as recently described (*9*, *10*). In brief, 5 μg/ml of VWF dimers in near-physiologic buffer were incubated on a poly-L-lysine-coated mica substrate for 30 s, which was subsequently rinsed with water and finally dried in a gentle stream of nitrogen. AFM images of 1 μm × 1 μm and 1024 × 1024 pixels were recorded in tapping mode in air, using an MFP-3D AFM (Asylum Research, Santa Barbara, CA) and cantilevers with silicon tips (AC160TS, Olympus, Japan), possessing a nominal spring constant of 26 N/m and a resonance frequency of approximately 300 kHz. Raw image data were processed using SPIP software (v6.5.1; Image Metrology, Denmark). Image processing involved plane correction (third order polynomial plane-fitting), line-wise flattening (according to the histogram alignment routine), and Gaussian smoothing.

#### Data analysis

All data analysis was carried out using custom-written Matlab scripts (Matlab v.R2015b; The MathWorks Inc., Natick, MA) incorporated into a custom Matlab GUI. We obtained tether extension *vs*. time by subtracting the *z*-position of the reference bead from the *z*-position of the protein-tethered bead. All traces shown and analyzed are the raw extension *vs*. time traces recorded at 58 Hz, used without any filtering or smoothing. For ddFLN4 measurements, only beads that in unfolding force plateaus repeatedly showed a double-step with a short-lived intermediate state were taken into account for further analysis. Similarly, for VWF measurements, only beads repeatedly exhibiting two steps of equal height corresponding to unfolding of the A2 domains in unfolding force plateaus were analyzed, unless otherwise noted. Unfolding and refolding behavior for ddFLN4 and VWF under the different reported buffer conditions were observed in at least 3 independently prepared flow cells in all cases.

To determine the position of steps, we employed the step-finding algorithm by Kerssemakers et al. (*65*), and the corresponding change in extension was determined as the difference between the average extensions of the adjacent 1000 frames recorded before and after the step, respectively (fewer frames were used if the 1000-frame interval contained another step). Extensions of folding and unfolding (sub)steps were histogrammed for each clamped force (1 nm binning for ddFLN4, and 3 nm and 2 nm binning for VWF A2 unfolding and refolding, respectively), and fitted with Gaussians. Error bars in figures report the FWHM of the fits, divided by the square root of the respective counts. The resulting force–extension profiles were fitted to the WLC model of polymer elasticity (an approximation to this model with less than 1% relative error was used for fitting (*66*)). In the case of ddFLN4, a fixed persistence length of 0.5 nm was used to enable direct comparison with results from an AFM study by Schwaiger et al. (*4*). In the case of VWF A2, both persistence length and contour length were free fit parameters.

To determine the unfolding or refolding rates *k*(*F*) at a given constant force *F*, the respective fraction of observed unfolding or refolding events as a function of time was fitted with the exponential expression 1 – *a* exp(−*kt*) + *b* (**Supplementary Fig. S4**), where the free parameters *a* and *b* can compensate for events that were missed due to the finite measurement time or due to the finite time of motor movement when setting the force. However, such missed events were rare and parameters *a* and *b* were close to 1 and 0, respectively. Error bars on rates in figures indicate 95% confidence bounds of fits. In the case of VWF, only events corresponding to steps with extensions ≤ 60 nm were taken into account to ensure that only A2 unfolding events–and not dissociation of the D4-mediated intermonomer interaction (see **Supplementary Fig. S6**)– are analyzed.

The force dependence of unfolding and refolding rates was described by a single barrier kinetic model: *k*(*F*) = *k*_0_ exp(*F*Δ*x*/*k*_B_*T*), with the rate at zero force *k*_0_ and the distance to the transition state Δ*x* as fit parameters. Fitting was carried out as linear fits to the natural logarithm of the data. Error margins for *k*_0_ and Δ*x* given in the text correspond to 1 SD.

For bead rupture measurements, lifetimes at different constant forces were determined from the survival fraction vs. time data based on > 35 rupture events for each condition. In the case of mSA-beads, data were described by a single-exponential decay, and the corresponding lifetime was determined by a linear fit to the natural logarithm of the data. In the case of the more complex decay behavior observed for commercial streptavidin-coated beads, lifetimes for the fastest-and slowest-decaying populations were estimated by linear fits to the natural logarithm of the first and last 20% of data points, respectively. The dependence of estimated lifetimes on force was again described by the single barrier kinetic model introduced above.

## Acknowledgements

### General

We thank Wolfgang Ott and Lukas F. Milles for providing ELP and ddFLN4 constructs, respectively, Gesa König and Thomas Nicolaus for laboratory assistance, and Ellis Durner, Reinhard Schneppenheim, and Hermann E. Gaub for helpful discussions.

### Funding

This study was supported by research funding from the German Research Foundation (DFG) to the projects “Unraveling the Mechano-Regulation of Von Willebrand Factor” (project number 386143268), “Shear Flow Regulation of Hemostasis - Bridging the Gap between Nanomechanics and Clinical Presentation” (SHENC, project FOR1543), SFB 863 project A11, and the Nanosystems Initiative Munich.

### Author contributions

A.L., P.U.W., S.M.S., M.A.B., M.B. and J.L. designed experiments. T.O. designed and prepared VWF constructs. S.M.S. prepared mSA constructs. A.L. prepared samples. A.L., P.U.W., and S.M.S. performed experiments. P.U.W. wrote analysis software. A.L. and P.U.W. analyzed results. A.L. prepared figures. A.L., M.B. and J.L. wrote the manuscript. All authors discussed the data and critically reviewed the manuscript.

### Competing interests

The authors declare no competing interests.

### Data and materials availability

All data presented in the main and supplementary figures are available from the authors upon request. The employed MT setup control and bead-tracking software, developed by Cnossen et al. (*15*), is open-source and available from http://www.github.com/jcnossen/BeadTracker. Custom-written Matlab software for analysis of magnetic tweezers data is available from the authors upon request. Plasmids for the ELP linkers used in this study are provided at Addgene by Ott et al. (Addgene accession number 90472).

## Supplementary Materials

### Supplementary Figures

**Fig. S1.**
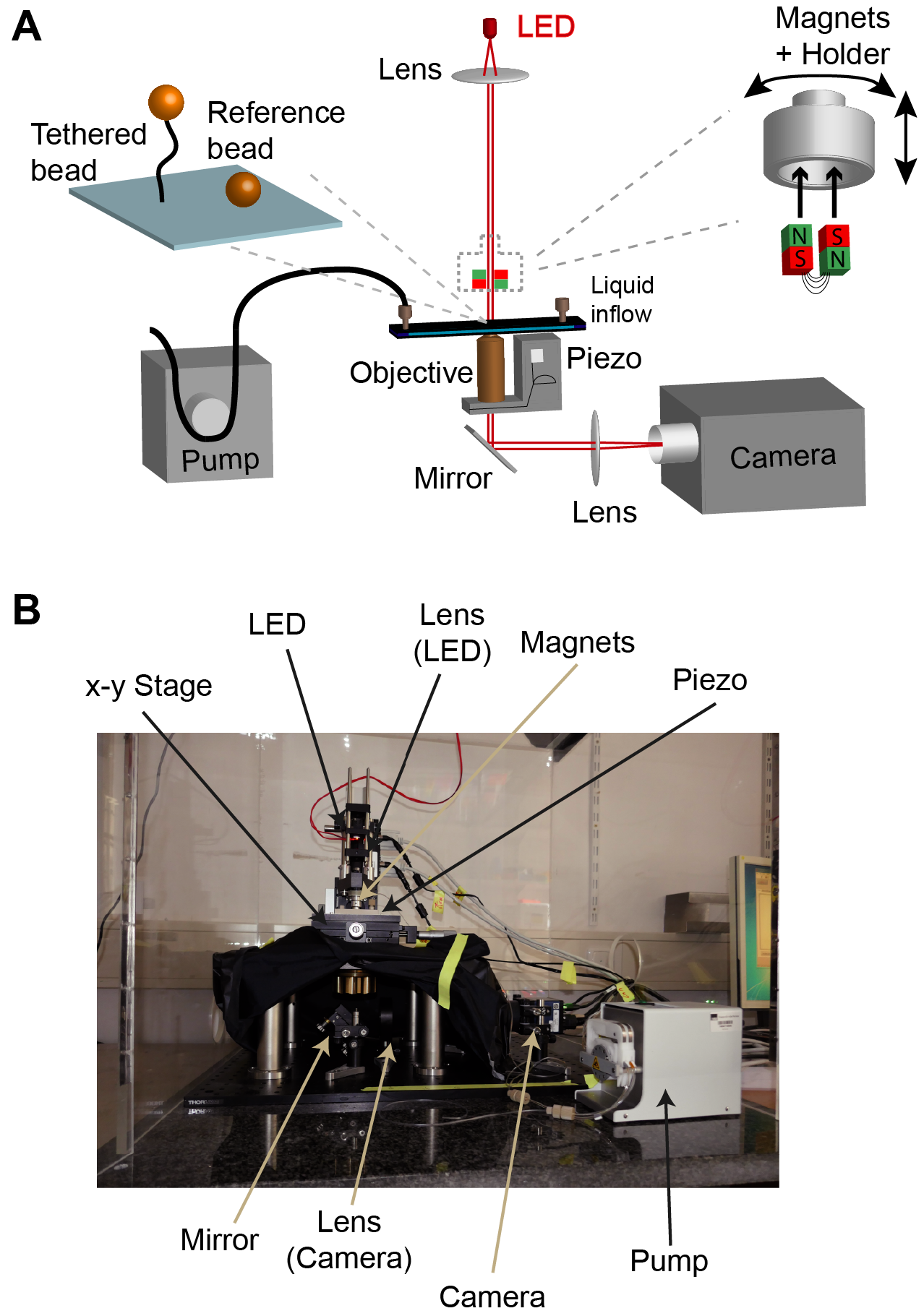
Magnetic tweezers setup. **(A)** Schematic of the MT setup. Proteins are tethered between a magnetic bead and the bottom surface of the flow cell, which is illuminated using an LED. A large field of view is imaged using a 40x oil-immersion objective and a CMOS sensor camera. For creating the look-up table necessary to track the *z* position of the beads, the objective is mounted on a piezo stage. A set of two cubic permanent magnets is positioned above the flow cell. The distance between magnets and flow cell can be adjusted using a DC-motor in order to adjust the force applied to the magnetic beads. A peristaltic pump allows for flushing the flow cell. For technical details of the different components, see **Methods**. **(B)** Image of the MT setup with essential components being highlighted.

**Fig. S2.**
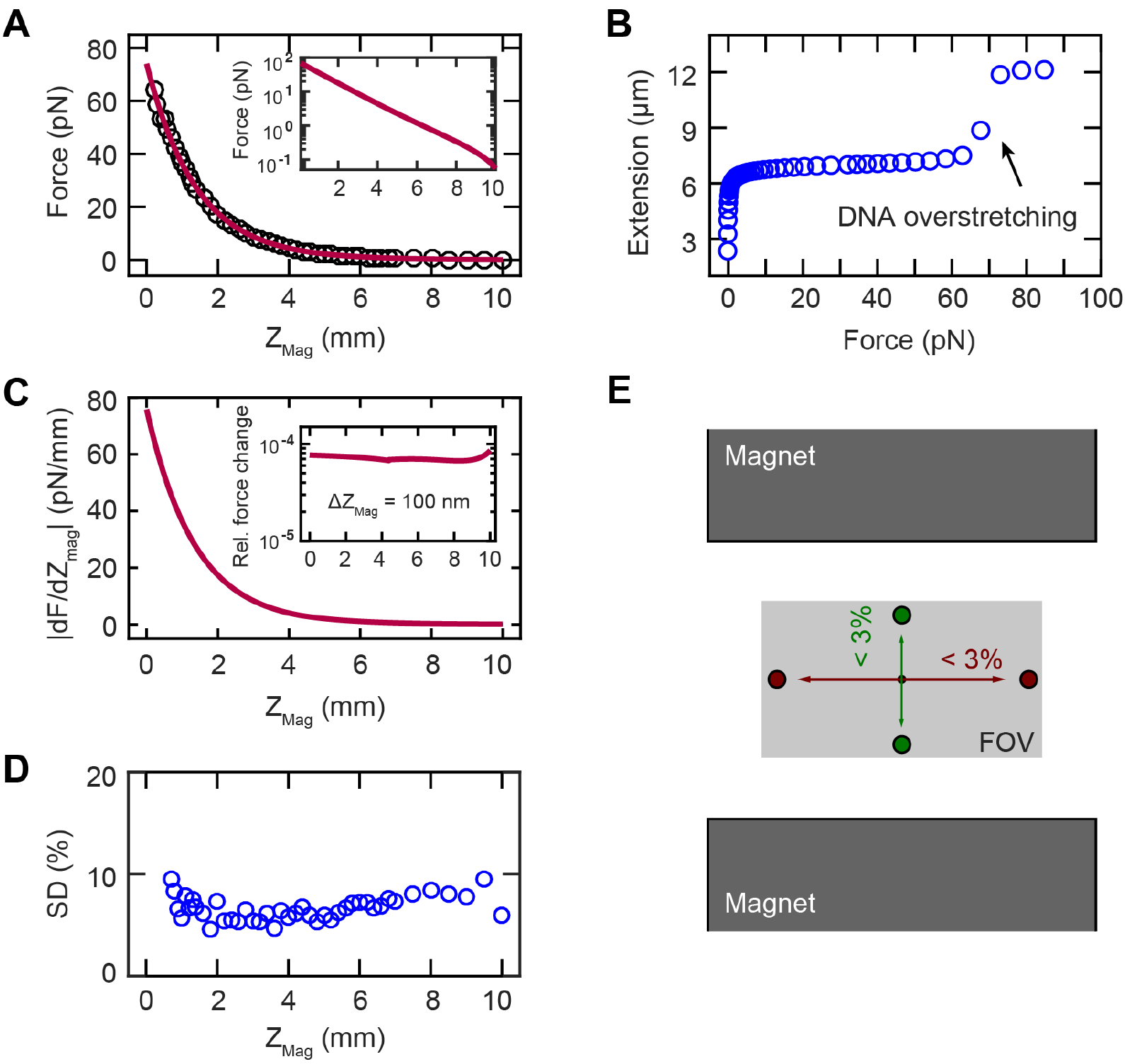
Force calibration of the MT setup. **(A)** Force acting on the magnetic beads used in this study (Dynabeads M270) as a function of the distance *Z*_Mag_ between the magnets and the flow cell. Forces were calibrated using the method described by Velthuis et al. (*62*), based on the Brownian fluctuations of long (here 21 kbp) double-stranded (ds) DNA tethers. Data points are mean forces determined from 16 DNA tethers. The green line is the final fit of the dependence of force on the magnet distance. **(B)** Exemplary trace of a 21 kbp dsDNA tether, showing the B-S overstretching transition at the expected force of ~65 pN, confirming the force calibration from analysis of the transverse fluctuations. **(C)** Absolute value of the derivative of the force with respect to *Z*_Mag_. The inset shows the relative force change for extension changes in *z* direction of 100 nm -larger than any (un-)folding steps in our measurements-, which was found to be < 10^−4^ for all forces, as calculated from the expression for |d*F*/dZ_Mag_|. **(D)** Bead-to-bead force variation. Independently performing the calibration procedure for 16 different DNA tethers, we found the standard deviation of the force from the mean value to be **≲** 10% over the whole range of magnet distances, indicating a bead-to-bead force variation of **≲** 10%, in line with previous reports (*14*, *63*, *64*). **(E)** Force uniformity across the field of view. To verify that the magnetic field is uniform and thus the forces do not vary significantly across the field of view (FOV), we repeatedly performed the force calibration procedure for the same DNA tether at different positions at the edges of the FOV, as schematically indicated by circles, and in the middle of the FOV. For each of four independently measured DNA tethers, changes in force were found to be < 3% both along the axis parallel to and the axis perpendicular to the gap between the magnets (not drawn to scale).

**Fig. S3.**
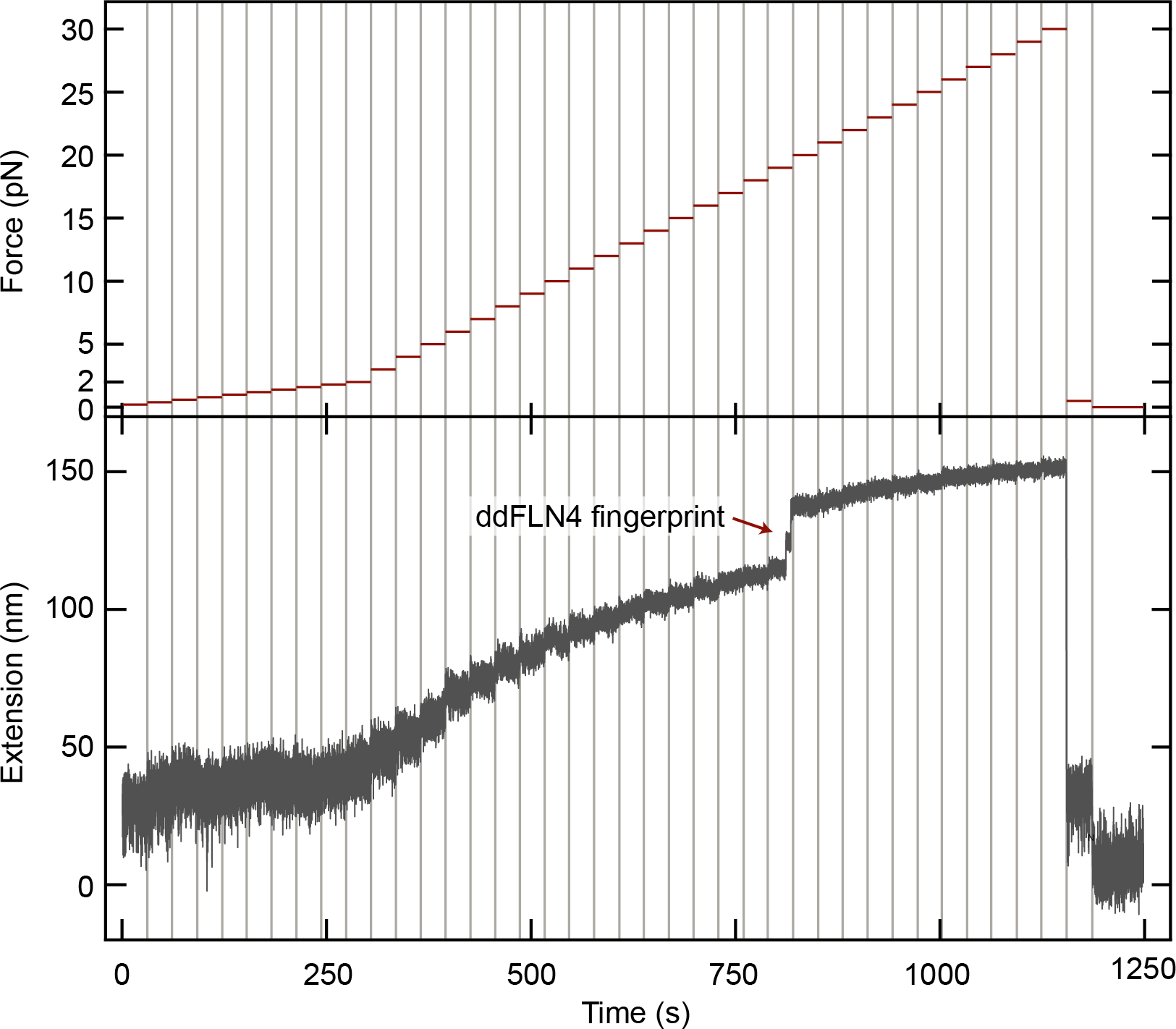
Extension of ELP linker-protein tethers. Exemplary extension trace (bottom) of a ddFLN4-ELP linker complex tethered between glass surface and magnetic bead as shown in Fig. 1 in the main text, recorded while the force was increased stepwise every 30 s (indicated by red lines; top), in steps of 0.2 pN between 0.2 and 2 pN, and in steps of 1 pN between 2 and 30 pN. Afterwards, the tether was relaxed to 0.5 pN to allow for refolding of ddFLN4 and further relaxed to zero force to determine the zero position of extension. No peculiar features-in particular no steps-were observed over the entire probed force range, with exception of the characteristic ddFLN4 unfolding pattern, which served to identify specific single-tethered beads. This finding shows that the ELP linker does not cause any signals that may interfere with analysis of the specific signals of the measured protein of interest.

**Fig. S4.**
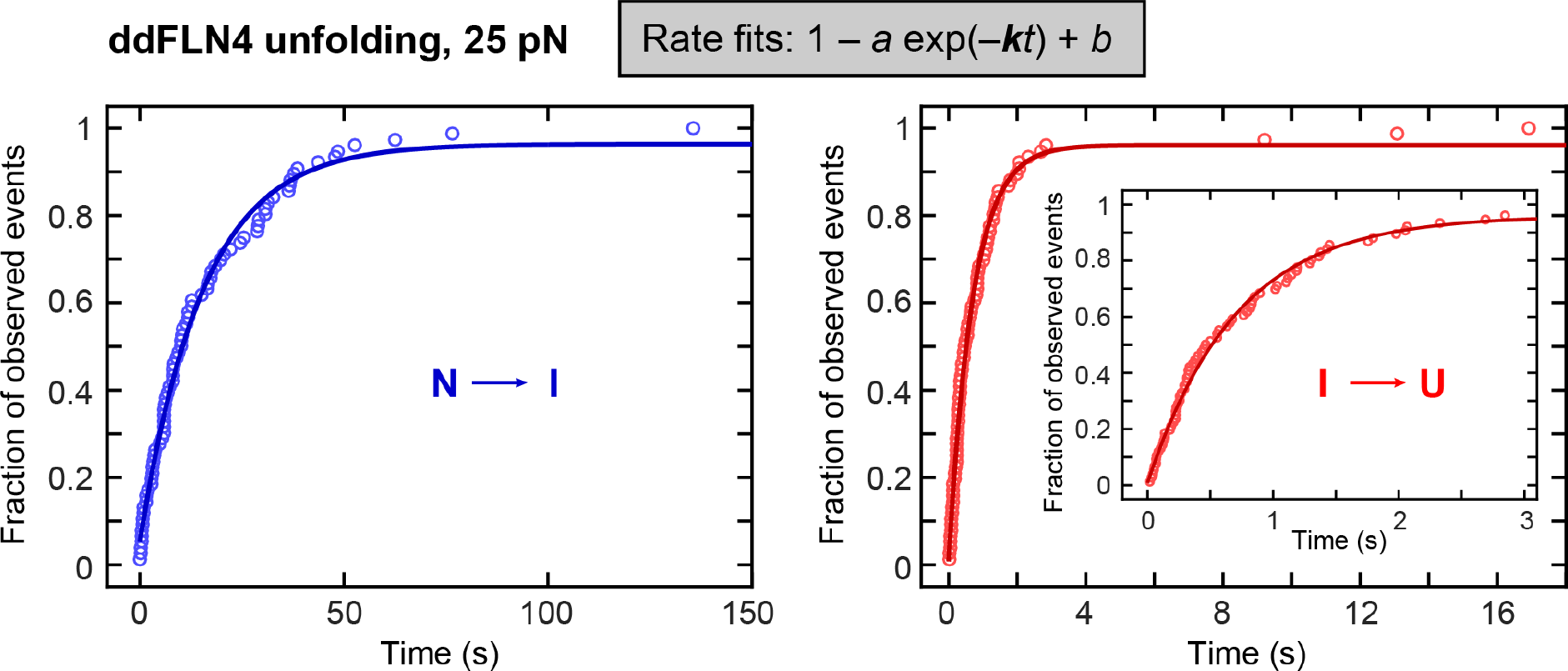
Determination of rates from the observed unfolding and refolding events. Concept of rate determination from the fraction of observed unfolding or refolding events as a function of time. Shown here as an example are the fractions of observed unfolding events *vs*. time for the two substeps of ddFLN4 unfolding at 25 pN, i.e. for the transitions from the native (N) to the intermediate (I) state (left, blue) and from the intermediate to the unfolded (U) state (right, red). To obtain the unfolding rate *k* of a transition at constant force *F*, the fraction of observed unfolding events as a function of time *t* is fit to the expression 1 - *a* exp(−*kt*) + *b* (lines), where the free parameters *a* and *b* can compensate for events that were missed due to the finite measurement time or due to the finite time of motor movement when setting the force. As a rule, parameters *a* and *b* were close to 1 and 0, respectively.

**Fig. S5.**
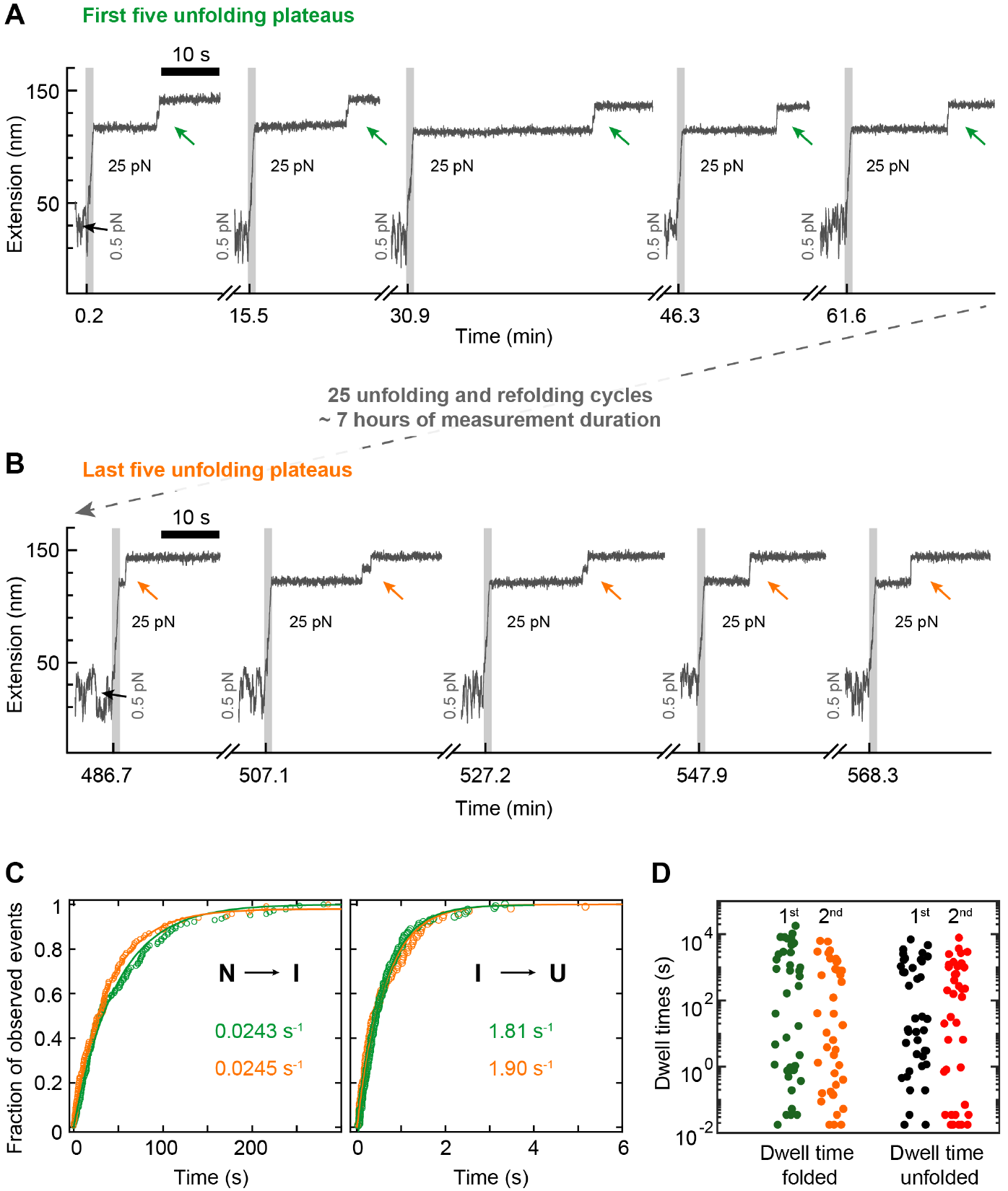
ddFLN4 does not exhibit hysteresis upon repeated unfolding and refolding. **(A,B)** Segments from a ~10 h-long measurement on ddFLN4 tethers with repeated alternating unfolding and refolding plateaus (similar to the data shown in **Fig. 2B**). Reliable unfolding and refolding was observed throughout the entire measurement. Shown here for one exemplary tether are the first five **(A)** and the last five **(B)** unfolding plateaus (all at 25 pN; unfolding events marked by arrows), which were separated by 25 cycles of unfolding and refolding, corresponding to ≈7 h of measurement duration. We analyzed the same 31 ddFLN4 tethers, separately for the first five and last five unfolding plateaus. The obtained mean extension values for the two unfolding transitions N→I and I→U both varied by less than 4%. Furthermore, the measured unfolding rates matched very closely. **(C)** Fits and unfolding rates are shown in green and orange for the first five and last five plateaus, respectively. Both for the first step of unfolding, N→I (left panel), and for the second step of unfolding, I→U (right panel), rates deviated by less than 5%, well within the 95% confidence intervals of the fits. Our data thus indicate that no significant hysteresis effects occur for ddFLN4 even after tens of unfolding/refolding cycles and spending an extended period of time in the unfolded state. **(D)** Analysis of a long (~50 h) trace at constant force close to the equilibrium point (the trace shown in **Fig. 3B**). The dwell times in the folded and unfolded states were quantified and are shown separately for the first and second halves of the trace. The distributions for the two halves of the trace for both folded and unfolded states are identical, within experimental error (as assessed by a two-sample Kolmogorov-Smirnov test with *p* = 0.51 and *p* = 0.53, respectively).

**Fig. S6.**
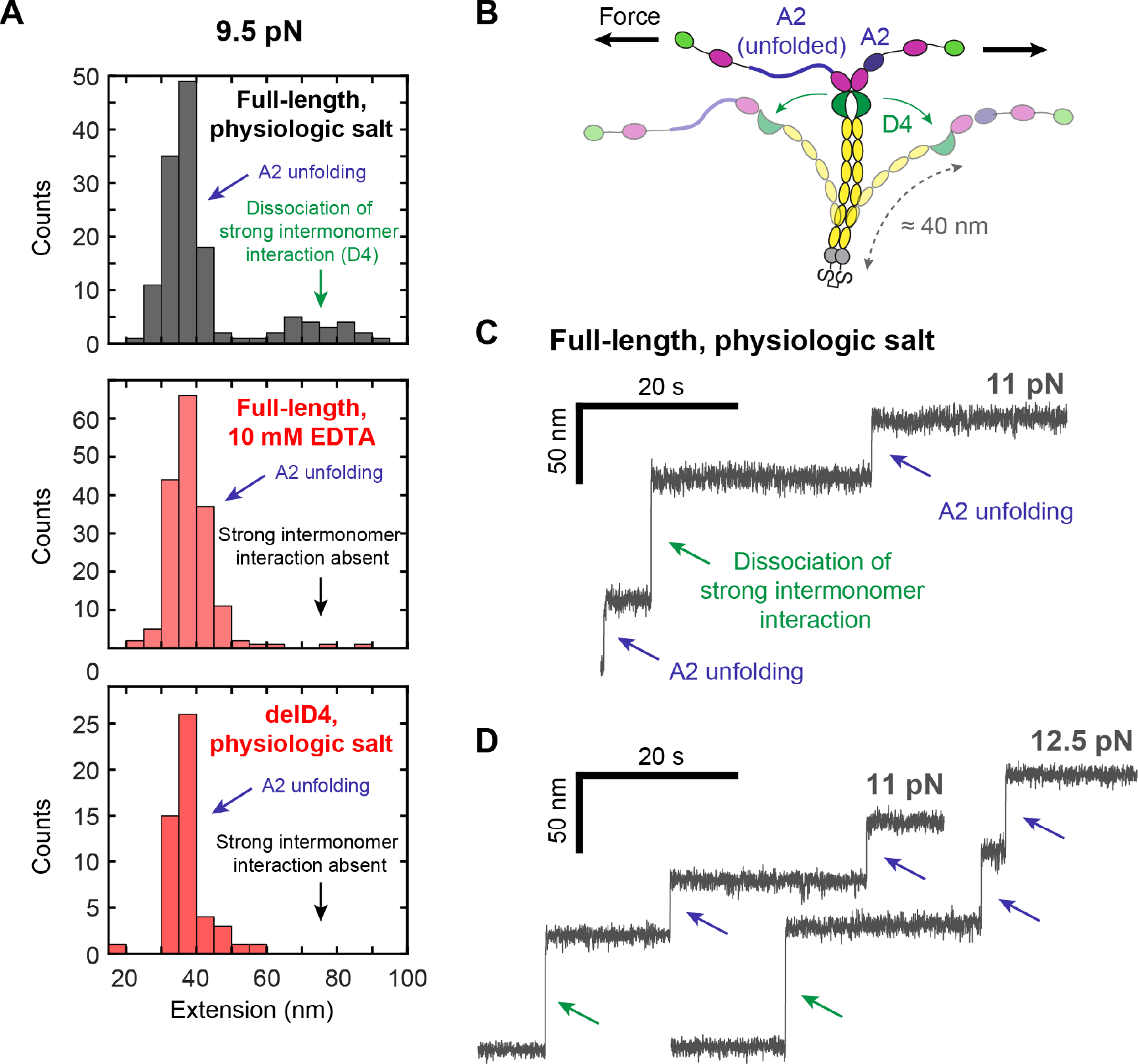
Dissociation of D4-mediated intermonomer interaction in VWF dimers. **(A)** Extension histograms of steps observed in traces of VWF dimers recorded at a force of 9.5 pN, for full-length dimers in the presence of divalent ions (top) or in the presence of 10 mM EDTA (middle), and for dimers with a deletion of the D4 domain (delD4, see also **Supplementary Fig. S7**) in the presence of divalent ions (bottom). In the case of the full-length dimers, in the presence of divalent ions a broad peak at extension values of roughly 7080 nm is observed in addition to the peak associated with A2 unfolding, centered at ca. 36 nm. In the presence of EDTA, or for the delD4 construct, in contrast, only the peak associated with A2 unfolding is observed. The length increase by 70-80 nm, the sensitivity to removal of divalent ions by EDTA, and the involvement of the D4 domain are in line with the dissociation of a strong intermonomer interaction mediated by VWF’s D4 domain that has recently been identified in AFM force measurements on VWF dimers (*9*, *10*). **(B)** Schematic of dimer opening. Dissociation of an intermonomer interaction mediated by the D4 domain (green) leads to the opening of the closed stem region of VWF (yellow) and thus a release of formerly hidden length of approximately 80 nm. Dimer opening occurs independently of A2 (blue) unfolding, since the A2 domains are not shielded from force by the D4-mediated interaction. **(C)** Exemplary extension trace of a full-length dimer exhibiting unfolding of both A2 domains and dimer opening, recorded at 11 pN. **(D)** Extension traces from the same VWF dimer tether, probed at different forces and repeatedly exhibiting dimer opening, implying reversibility of the D4-mediated intermonomer interaction.

**Fig. S7.**
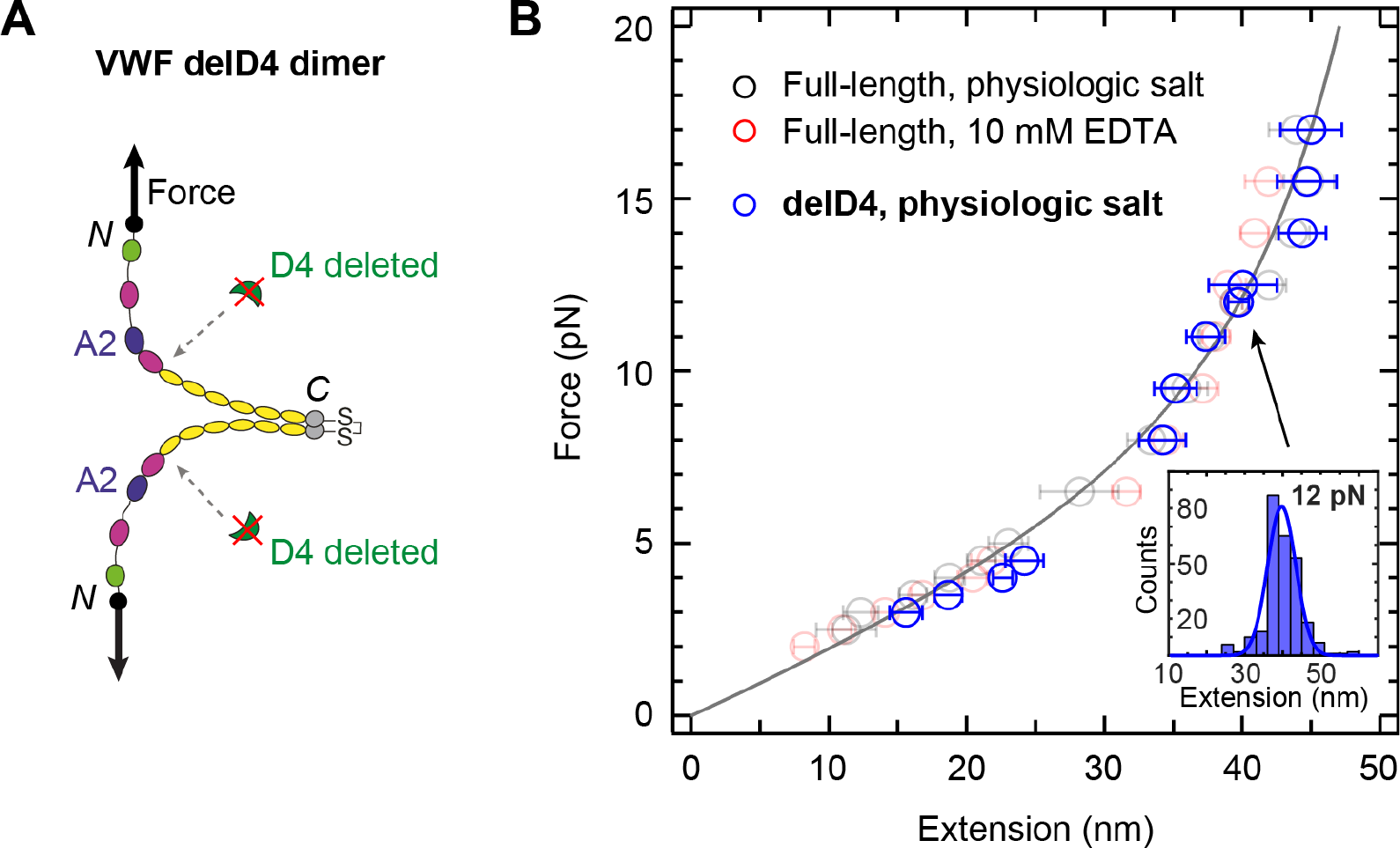
Measurements on VWF dimers with deletion of the D4 domain. **(A)** Schematic structure of a VWF dimer with deletion of both D4 domains (delD4 dimer). The two A2 domains are shown in blue. Arrows indicate the direction of force acting on the two N termini during MT experiments. **(B)** Force-extension profile of A2 unfolding and refolding, recorded for the delD4 construct in near-physiologic buffer at pH 7.4 (blue symbols). The force-extension profile closely matches those obtained for the full-length construct in near-physiologic buffer and in buffer with 10 mM EDTA (co-plotted with lower opacity in black and red, respectively), as presented in Fig. 4c in the main text. The line is the global WLC fit to all data from the full-length construct, as presented in the inset in Fig. 4c in the main text. Data points are obtained by Gaussian fits to step extension histograms (inset) at each constant force. Data points above 5 pN are from unfolding, data points up to 5 pN from refolding. Error bars correspond to the FWHM of Gaussian fits, divided by the square root of counts.

**Fig. S8.**
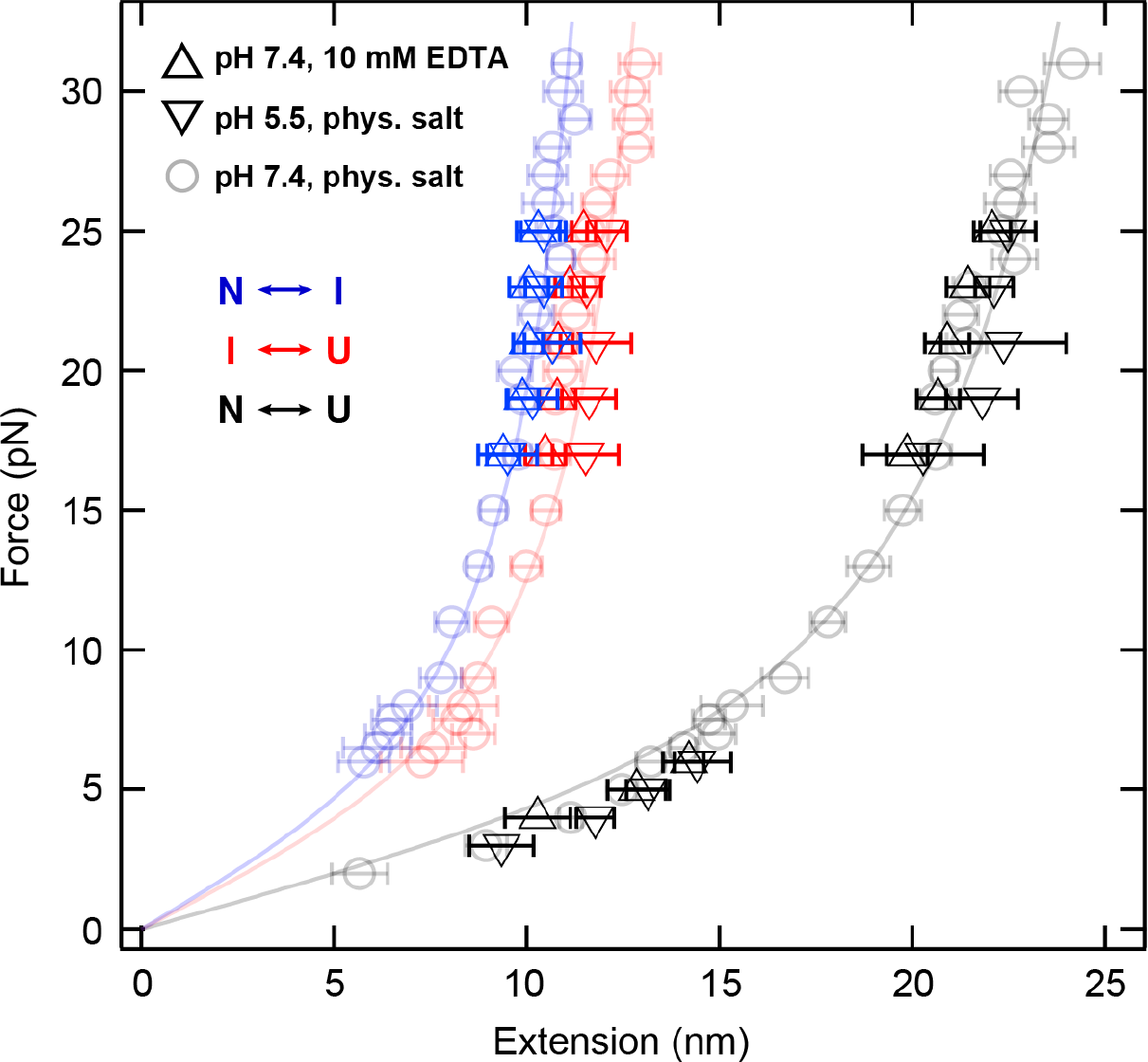
ddFLN4 unfolding and refolding under varied salt and pH conditions. Extension of ddFLN4 unfolding and refolding steps at different constant forces and under varied buffer conditions. Extensions of the transitions between the native state and the intermediate state (blue) as well as between the intermediate and the unfolded state (red) are shown separately in addition to the full extension between native and unfolded state (black). Data points at forces up to 8 pN are from refolding, data points at forces above 8 pN from unfolding measurements. Co-plotted with lower opacity are the data obtained for near-physiological buffer conditions (pH 7.4, with divalent ions; circles) as shown in Fig. 2c in the main text and the respective WLC fits (lines). Force-extension data sets obtained at pH 7.4 in the presence of 10 mM EDTA (upward triangles) and at acidic pH 5.5 in the presence of divalent ions (downward triangles) both are within measurement uncertainty identical to the ones obtained for near-physiologic buffer conditions. Error bars correspond to the FWHM of Gaussian fits, divided by the square root of counts.

**Fig. S9.**
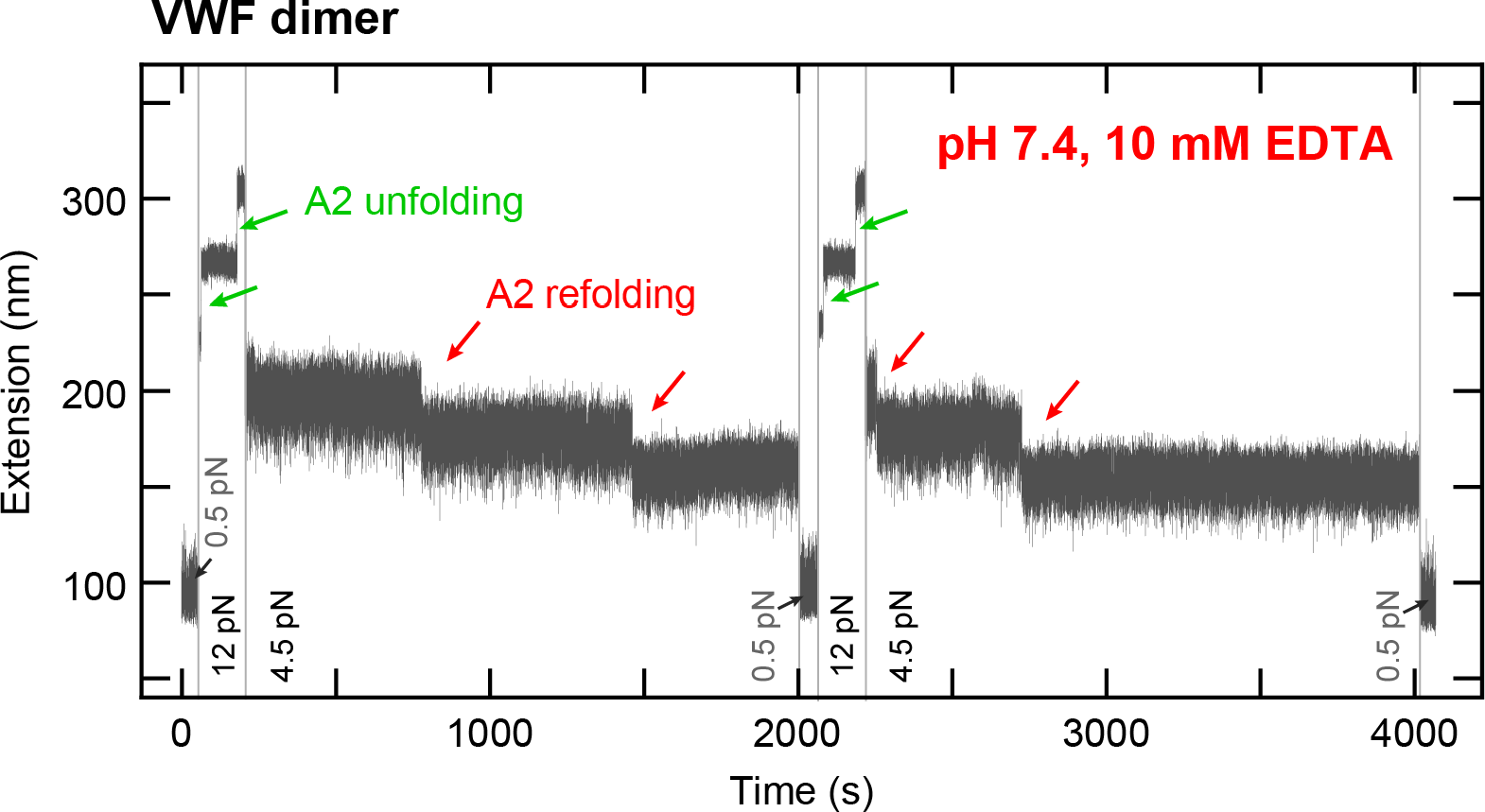
Refolding of the VWF A2 domain under mechanical load in the absence of Ca^2+^. Extension *vs*. time trace of a VWF dimer tether subjected to alternating intervals of high force (here 12 pN), allowing for A2 unfolding, of intermediate force (here 4.5 pN), allowing for direct observation of A2 refolding, and of low force (0.5 pN) to ensure refolding, in buffer without Ca^2+^ and with 10 mM EDTA. Unfolding and refolding of the two A2 domains are observed as two independent positive or negative steps in the trace, respectively. Direct observation of refolding steps (marked by red arrows) shows that A2 can refold under significant mechanical load even in the absence of Ca^2+^.

**Fig. S10.**
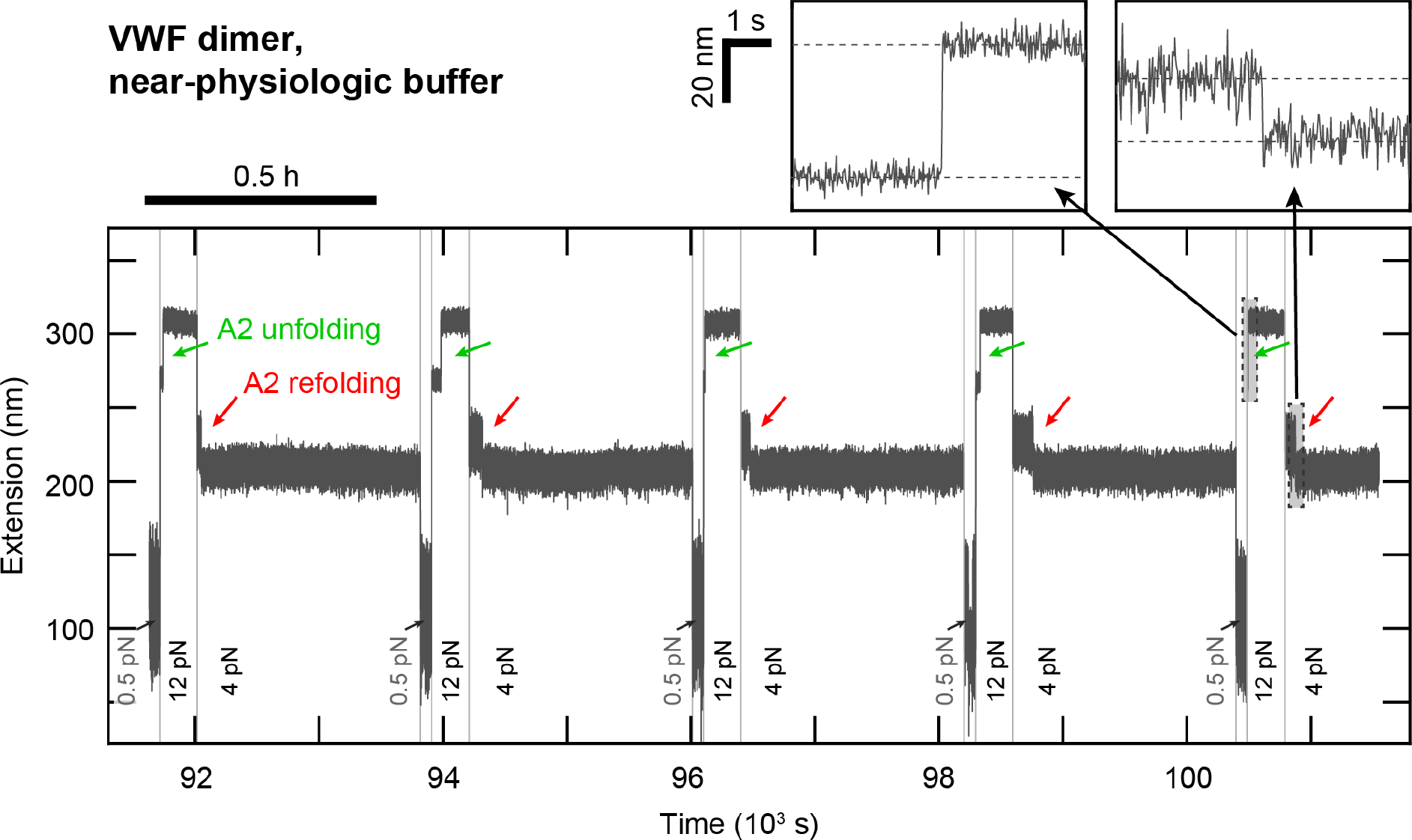
Occasional inhibition of A2 refolding in VWF dimer tethers. Segment of a ≈30 h-long extension *vs*. time trace of a VWF dimer tether subjected to alternating intervals of high force (here 12 pN), allowing for A2 unfolding, and of different intermediate forces (4 pN shown here), allowing for direct observation of A2 refolding, recorded under near-physiologic buffer conditions. The shown tether exhibits the unfolding and refolding signal (marked by arrows) of only one of the two A2 domains. Otherwise, the tether does not show any differences to regular tethers exhibiting signals of both A2 domains. In particular, the observed A2 unfolding and refolding steps were indistinguishable (see insets). In such tethers exhibiting only one A2 signal, which occurred only occasionally, refolding of one of the A2 domains may be inhibited due to cis-trans isomerization of a cis-proline, as reported in a previous OT study (*8*).

## References

1. E. M. Puchner, H. E. Gaub, Single-Molecule Mechanoenzymatics. Annu. Rev. Biophys. 41, 497–518 (2012).

2. V. Vogel, Mechanotransduction involving multimodular proteins: converting force into biochemical signals. Annu. Rev. Biophys. Biomol. Struct. 35, 459–88 (2006).

3. I. Schwaiger, A. Kardinal, M. Schleicher, A. A. Noegel, M. Rief, A mechanical unfolding intermediate in an actin-crosslinking protein. Nat. Struct. Mol. Biol. 11, 81–5 (2004).

4. I. Schwaiger, M. Schleicher, A. A. Noegel, M. Rief, The folding pathway of a fast-folding immunoglobulin domain revealed by single-molecule mechanical experiments. EMBO Rep. 6, 46–51 (2005).

5. T. A. Springer, Von Willebrand factor, Jedi knight of the bloodstream. Blood. 124, 1412–1426 (2014).

6. A. Löf, J. P. Müller, M. A. Brehm, A biophysical view on von Willebrand factor activation. J. Cell. Physiol. 233, 799–810 (2018).

7. H. Fu et al., Flow-induced elongation of von Willebrand factor precedes tension-dependent activation. Nat. Commun. 8, 324 (2017).

8. X. Zhang, K. Halvorsen, C.-Z. Zhang, W. P. Wong, T. A. Springer, Mechanoenzymatic Cleavage of the Ultralarge Vascular Protein von Willebrand Factor. Science. 324, 1330–1334 (2009).

9. J. P. Müller et al., Force Sensing by the Vascular Protein von Willebrand Factor is Tuned by a Strong Intermonomer Interaction. Proc. Natl. Acad. Sci. 113, 1208–1213 (2016).

10. J. P. Müller et al., pH-dependent interactions in dimers govern the mechanics and structure of von Willebrand Factor. Biophys. J. 111, 312–322 (2016).

11. K. C. Neuman, A. Nagy, Single-molecule force spectroscopy: optical tweezers, magnetic tweezers and atomic force microscopy. Nat. Methods. 5, 491–505 (2008).

12. S. B. Smith, L. Finzi, C. Bustamante, Direct mechanical measurements of the elasticity of single DNA molecules by using magnetic beads. Science. 258, 1122–6 (1992).

13. T. R. Strick, J. F. Allemand, D. Bensimon, A. Bensimon, V. Croquette, The elasticity of a single supercoiled DNA molecule. Science. 271, 1835–7 (1996).

14. J. Lipfert, X. Hao, N. H. Dekker, Quantitative modeling and optimization of magnetic tweezers. Biophys. J. 96, 5040–9 (2009).

15. J. P. Cnossen, D. Dulin, N. H. Dekker, An optimized software framework for real-time, high-throughput tracking of spherical beads. Rev. Sci. Instrum. 85, 103712 (2014).

16. N. Ribeck, O. A. Saleh, Multiplexed single-molecule measurements with magnetic tweezers. Rev. Sci. Instrum. 79, 094301 (2008).

17. I. De Vlaminck et al., Highly Parallel Magnetic Tweezers by Targeted DNA Tethering. Nano Lett. 11, 5489–5493 (2011).

18. B. M. Lansdorp, S. J. Tabrizi, A. Dittmore, O. A. Saleh, A high-speed magnetic tweezer beyond 10,000 frames per second. Rev. Sci. Instrum. 84, 044301 (2013).

19. D. Dulin et al., High Spatiotemporal-Resolution Magnetic Tweezers: Calibration and Applications for DNA Dynamics. Biophys. J. 109, 2113–25 (2015).

20. A. Huhle et al., Camera-based three-dimensional real-time particle tracking at kHz rates and Ångström accuracy. Nat. Commun. 6, 5885 (2015).

21. H. Chen et al., Dynamics of Equilibrium Folding and Unfolding Transitions of Titin Immunoglobulin Domain under Constant Forces. J. Am. Chem. Soc. 137, 3540–3546 (2015).

22. S. Haldar, R. Tapia-Rojo, E. C. Eckels, J. Valle-Orero, J. M. Fernandez, Trigger factor chaperone acts as a mechanical foldase. Nat. Commun. 8, 668 (2017).

23. I. Popa et al., A HaloTag Anchored Ruler for Week-Long Studies of Protein Dynamics. J. Am. Chem. Soc. 138, 10546–10553 (2016).

24. D. Spadaro et al., Tension-Dependent Stretching Activates ZO-1 to Control the Junctional Localization of Its Interactors. Curr. Biol. 27, 3783–3795.e8 (2017).

25. A. S. Adhikari, E. Glassey, A. R. Dunn, Conformational dynamics accompanying the proteolytic degradation of trimeric collagen I by collagenases. J. Am. Chem. Soc. 134, 1325965 (2012).

26. X. J. A. Janssen et al., Torsion stiffness of a protein pair determined by magnetic particles. Biophys. J. 100, 2262–7 (2011).

27. E. Pérez-Ruiz et al., Probing the Force-Induced Dissociation of Aptamer-Protein Complexes. Anal. Chem. 86, 3084–3091 (2014).

28. A. S. Adhikari, J. Chai, A. R. Dunn, Mechanical Load Induces a 100-Fold Increase in the Rate of Collagen Proteolysis by MMP-1. J. Am. Chem. Soc. 133, 1686–1689 (2011).

29. A. van Reenen, F. Gutiérrez-Mejía, L. J. van IJzendoorn, M. W. J. Prins, Torsion Profiling of Proteins Using Magnetic Particles. Biophys. J. 104, 1073–1080 (2013).

30. H. Chen et al., Mechanical perturbation of filamin A immunoglobulin repeats 20-21 reveals potential non-equilibrium mechanochemical partner binding function. Sci. Rep. 3, 1642 (2013).

31. M. Yao et al., Force-dependent conformational switch of α-catenin controls vinculin binding. Nat. Commun. 5, 4525 (2014).

32. M. Yao et al., The mechanical response of talin. Nat. Commun. 7, 11966 (2016).

33. S. Le et al., Mechanotransmission and Mechanosensing of Human alpha-Actinin 1. Cell Rep. 21, 2714–2723 (2017).

34. W. Ott et al., Elastin-like Polypeptide Linkers for Single-Molecule Force Spectroscopy. ACS Nano. 11, 6346–6354 (2017).

35. K. Halvorsen, W. P. Wong, Massively parallel single-molecule manipulation using centrifugal force. Biophys. J. 98, L53–5 (2010).

36. D. Yang, A. Ward, K. Halvorsen, W. P. Wong, Multiplexed single-molecule force spectroscopy using a centrifuge. Nat. Commun. 7, 11026 (2016).

37. G. Sitters et al., Acoustic force spectroscopy. Nat. Methods. 12, 47–50 (2015).

38. W. Ott, T. Nicolaus, H. E. Gaub, M. A. Nash, Sequence-Independent Cloning and Post-Translational Modification of Repetitive Protein Polymers through Sortase and Sfp-Mediated Enzymatic Ligation. Biomacromolecules. 17, 1330–1338 (2016).

39. C. S. Theile et al., Site-specific N-terminal labeling of proteins using sortase-mediated reactions. Nat. Protoc. 8, 1800–7 (2013).

40. J. Yin et al., Genetically encoded short peptide tag for versatile protein labeling by Sfp phosphopantetheinyl transferase. Proc. Natl. Acad. Sci. 102, 15815–15820 (2005).

41. M. W.-L. Popp, H. L. Ploegh, Making and Breaking Peptide Bonds: Protein Engineering Using Sortase. Angew. Chemie Int. Ed. 50, 5024–5032 (2011).

42. W. Ott, M. A. Jobst, C. Schoeler, H. E. Gaub, M. A. Nash, Single-molecule force spectroscopy on polyproteins and receptor-ligand complexes: The current toolbox. J. Struct. Biol. 197, 3–12 (2017).

43. E. Durner, W. Ott, M. A. Nash, H. E. Gaub, Post-Translational Sortase-Mediated Attachment of High-Strength Force Spectroscopy Handles. ACS Omega. 2, 3064–3069 (2017).

44. L. F. Milles, K. Schulten, H. E. Gaub, R. C. Bernardi, Molecular mechanism of extreme mechanostability in a pathogen adhesin. Science. 359, 1527–1533 (2018).

45. L. F. Milles, E. A. Bayer, M. A. Nash, H. E. Gaub, Mechanical Stability of a High-Affinity Toxin Anchor from the Pathogen Clostridium perfringens. J. Phys. Chem. B. 121, 3620–3625 (2017).

46. S. M. Sedlak et al., Monodisperse measurement of the biotin-streptavidin interaction strength in a well-defined pulling geometry. PLoS One. 12, e0188722 (2017).

47. E. Evans, K. Ritchie, Dynamic strength of molecular adhesion bonds. Biophys. J. 72, 1541–1555 (1997).

48. Z. T. Yew, M. Schlierf, M. Rief, E. Paci, Direct evidence of the multidimensionality of the free-energy landscapes of proteins revealed by mechanical probes. Phys. Rev. E. 81, 031923 (2010).

49. S. M. Sedlak et al., Nano Lett., in press, doi:10.1021/acs.nanolett.8b04045.

50. Y. F. Zhou et al., Sequence and structure relationships within von Willebrand factor. Blood. 120, 449–458 (2012).

51. A. J. Xu, T. A. Springer, Calcium stabilizes the von Willebrand factor A2 domain by promoting refolding. Proc. Natl. Acad. Sci. 109, 3742–3747 (2012).

52. C. Baldauf et al., Shear-induced unfolding activates von Willebrand factor A2 domain for proteolysis. J. Thromb. Haemost. 7, 2096–2105 (2009).

53. A. J. Jakobi, A. Mashaghi, S. J. Tans, E. G. Huizinga, Calcium modulates force sensing by the von Willebrand factor A2 domain. Nat. Commun. 2, 385 (2011).

54. M. Zhou et al., A novel calcium-binding site of von Willebrand factor A2 domain regulates its cleavage by ADAMTS13. Blood. 117, 4623–31 (2011).

55. O. K. Dudko, G. Hummer, A. Szabo, Theory, analysis, and interpretation of single-molecule force spectroscopy experiments. Proc. Natl. Acad. Sci. 105, 15755–15760 (2008).

56. Y.-F. Zhou et al., A pH-regulated dimeric bouquet in the structure of von Willebrand factor. EMBO J. 30, 4098–111 (2011).

57. S. Lippok et al., Shear-Induced Unfolding and Enzymatic Cleavage of Full-Length VWF Multimers. Biophys. J. 110, 545–554 (2016).

58. D. G. Gibson et al., Enzymatic assembly of DNA molecules up to several hundred kilobases. Nat. Methods. 6, 343–345 (2009).

59. T. G. Schmidt, A. Skerra, The Strep-tag system for one-step purification and high-affinity detection or capturing of proteins. Nat. Protoc. 2, 1528–1535 (2007).

60. J. Phan et al., Structural basis for the substrate specificity of tobacco etch virus protease. J. Biol. Chem. 277, 50564–72 (2002).

61. P. U. Walker, W. Vanderlinden, J. Lipfert, Dynamics and energy landscape of DNA plectoneme nucleation. Phys. Rev. E. 98, 042412 (2018).

62. A. J. W. Te Velthuis, J. W. J. Kerssemakers, J. Lipfert, N. H. Dekker, Quantitative guidelines for force calibration through spectral analysis of magnetic tweezers data. Biophys. J. 99, 1292–302 (2010).

63. I. De Vlaminck, T. Henighan, M. T. J. van Loenhout, D. R. Burnham, C. Dekker, Magnetic Forces and DNA Mechanics in Multiplexed Magnetic Tweezers. PLoS One. 7, e41432 (2012).

64. E. Ostrofet, F. S. Papini, D. Dulin, Correction-free force calibration for magnetic tweezers experiments. Sci. Rep. 8, 15920 (2018).

65. J. W. J. Kerssemakers et al., Assembly dynamics of microtubules at molecular resolution. Nature. 442, 709–712 (2006).

66. R. Petrosyan, Improved approximations for some polymer extension models. Rheol. Acta. 56, 21–26 (2017).

67. P. Fucini, C. Renner, C. Herberhold, A. A. Noegel, T. A. Holak, The repeating segments of the F-actin cross-linking gelation factor (ABP-120) have an immunoglobulin-like fold. Nat. Struct. Biol. 4, 223–30 (1997).

68. W. Humphrey, A. Dalke, K. Schulten, VMD: visual molecular dynamics. J. Mol. Graph. 14, 33-8, 27–8 (1996).

